# Attenuation of IFITM proteins’ antiviral activity through sequestration into intraluminal vesicles of late endosomes

**DOI:** 10.1101/2025.05.27.656272

**Authors:** David Prikryl, You Zhang, Gregory B. Melikyan

## Abstract

Interferon-induced transmembrane proteins (IFITMs) inhibit the entry of diverse enveloped viruses. The spectrum of antiviral activity of IFITMs is largely determined by their subcellular localization. IFITM1 localizes to and primarily blocks viral fusion at the plasma membrane, while IFITM3 prevents viral fusion in late endosomes by accumulating in these compartments. We have previously reported that cyclosporine A (CsA) treatment relieves the fusion block for the Influenza A virus, likely by relocating IFITM1 and IFITM3 from the plasma membrane and endosomes, respectively, to the Golgi area. Here, we report the existence of at least two distinct pools of IFITMs in CsA treated cells. While immunostaining of CsA treated cells using mild permeabilization agents, such as digitonin, suggests preferential IFITM localization at the Golgi apparatus, a harsher permeabilization protocol reveals a large, previously unidentified pool of IFITMs in late endosomes. Notably, IFITM redistribution was not associated with its degradation. A disproportionate loss of antibody access to the cytoplasmic N-terminus compared to the extracellular C-terminus of IFITMs after CsA treatment is consistent with sequestration of the N-terminal domain inside intraluminal vesicles of late endosomes. Accordingly, super-resolution microscopy reveals that CsA induces IFITM3 redistribution from the periphery to the interior of late endosomes. Together, our results imply that IFITMs relocate to intraluminal vesicles of late endosomes in the presence of CsA, thereby enabling viral fusion with the limiting membrane of these compartments. Our findings highlight the critical role of IFITM trafficking in antiviral defense and suggest a novel mechanism through which CsA modulates the cell’s susceptibility to viral infections.

## Introduction

Interferon-induced transmembrane proteins (IFITMs) impose a barrier to fusion of diverse enveloped viruses, such as the Influenza A Virus (IAV), Vesicular Stomatitis Virus (VSV), Respiratory syncytial virus. Dengue Virus, Ebola Virus, Measles Virus and other pathogenic viruses. Notable exceptions include the Murine Leukemia Virus and arenaviruses, such as the Lassa Virus (LASV), which are insensitive to IFITM restriction [1–4]. The human genome encodes for five IFITM proteins, with IFITM1, IFITM2, and IFITM3 exhibiting antiviral activity [5–7]. The significance of IFITM-mediated virus restriction *in vivo* is underscored by studies demonstrating that *Ifitm3* knockout mice succumb to IAV or Respiratory syncytial virus infection [8–11]. Additionally, several groups have established a correlation between single-nucleotide polymorphisms (SNPs) in the *Ifitm3* gene and more severe outcomes of IAV or Severe acute respiratory syndrome coronavirus 2 (SARS-CoV-2) infection [12–16].

The range of restricted viruses is largely determined by subcellular localization of IFITMs, which is regulated predominantly by the YXXL endocytic sorting motif within the N-terminal domain of IFITM2 and IFITM3. This motif is recognized by the clathrin adaptor protein 2 (AP2), which drives their internalization and concentration in endosomes, effectively preventing viruses from entering cells through an endocytic route [17–20]. Conversely, IFITM1 lacking the N-terminal endocytic signal predominantly resides in the plasma membrane (PM) and is more efficacious against viruses that tend to fuse at this location [2,5,21,22]. However, this rule is not without exceptions, since IFITM1 more potently hinders the entry and replication of filoviruses, such as Ebola virus, in some cell lines [3].

The IFITMs’ antiviral activity is further modulated through various post-translational modifications, including S-palmitoylation, ubiquitylation, phosphorylation, and methylation [23–28]. Subcellular localization and antiviral activity of IFITMs can be also altered by treatment with certain compounds, as reported by us and other groups [29–32]. However, the results on changes in subcellular localization of IFITM3 in treated cells differ between groups. We have found that cyclosporine A (CsA) triggered a rapid relocalization of IFITMs to the Golgi area without a noticeable degradation of these proteins. In contrast, others reported a strong colocalization of IFITM3 with the endolysosomal compartments that promote degradation after prolonged treatment with either rapamycin or cyclosporine H [29–31]. However, the mechanism by which cyclosporines modulate the localization and abundance of IFITMs remains unclear.

Here, to address the above discrepancies regarding the mechanism of cyclosporine antagonism with IFITMs’ function, we carried out comprehensive studies of CsA-driven IFITMs relocalization under varying conditions and correlated them with rescue of IAV fusion. Our results imply that CsA redirects IFITM1 and IFITM3 from the PM and the limiting membrane of endosomes, respectively, to the intraluminal vesicles (ILVs) of late endosomes. Such massive relocalization is not detectable in mildly permeabilized cells due to poor ILV accessibility to antibodies. We confirmed the CsA-induced IFITM3 redistribution to ILVs by super-resolution microscopy. These findings reconcile the reported differences in protein distribution and abundance and provide a plausible mechanism of CsA-mediated rescue of viral fusion in IFITM-expressing cells. This mechanism involves an effective removal of IFITMs from the PM and the limiting membrane of late endosomes where productive viral fusion takes place.

## Results

### CsA treatment renders a large pool of IFITMs inaccessible to antibodies in mildly permeabilized cells

We have previously shown that pretreatment of A549 cells ectopically expressing IFITM3 with a combination of cycloheximide (CHX) and CsA rescues IAV fusion with these cells without significantly reducing the level of IFITM3 assessed by Western blotting [32]. Inhibition of IFITM3 synthesis with CHX alone cleared the Golgi from the newly synthesized protein but did not alter the peripheral IFITM3 signal associated with the intracellular vesicles and the plasma membrane. Surprisingly, a combination of CsA and CHX caused a marked loss of IFITM3 signal in immunofluorescence experiments (Fig. 1A, B), in stark contrast with the Western blotting results, showing no IFITM3 degradation under these conditions (Fig. 1C and [32]). Note that treatment with CsA alone resulted in a significant loss of IFITM3 signal in immunostained samples.

**Figure 1.**
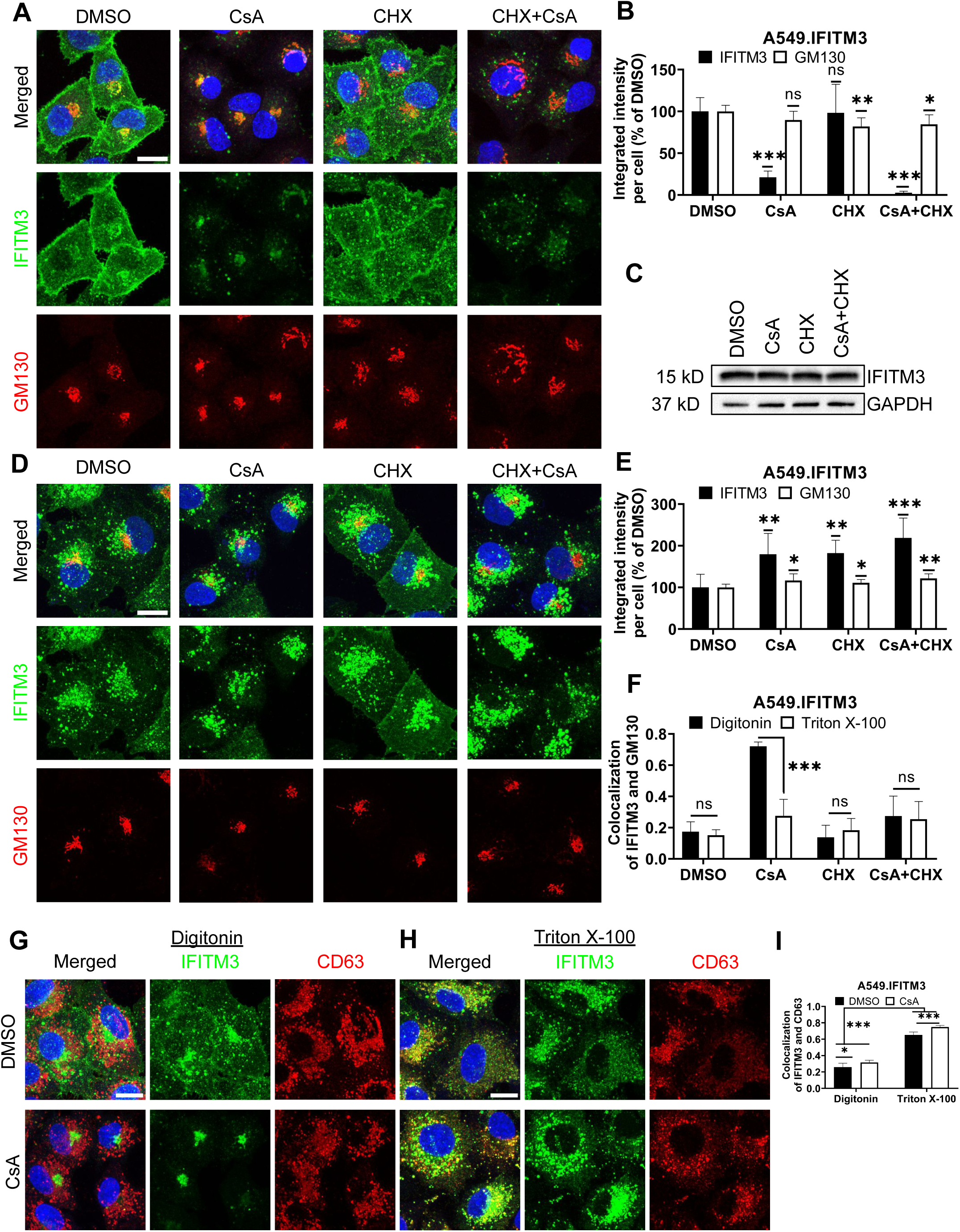
CsA treatment limits antibody access to IFITMs in digitonin-permeabilized cells. (A) A549.IFITM3 cells were treated with DMSO, CsA (20 µM), CHX (10 µg/mL), or a combination of CsA and CHX for 90 minutes, fixed, permeabilized with digitonin, and stained with anti-N-terminus of IFITM3 and anti-GM130 antibodies. (B) The integrated intensity of both signals per cell was measured and normalized to DMSO control. (C) Cells were treated as in (A), harvested, and cell lysates were analyzed by Western blotting. (D) A549.IFITM3 cells were treated as in (A), fixed, permeabilized with TX-100, and stained for IFITM3 and GM130. (E) Integrated intensities of IFITM3 and GM130 per cell normalized to DMSO control were calculated. (F) Colocalization of IFITM3 and GM130 signals was measured by calculating the Pearson’s coefficient. (G, H) A549.IFITM3 cells were treated either with DMSO or CsA (20 µM) for 90 minutes, fixed, permeabilized with either digitonin (G) or TX-100 (H), and immunostained for IFITM3 and CD63. Scale bars in A, D and G are 10 µm. (I) Colocalization of IFITM3 and CD63 signals was calculated as in (F). Data are means and S.D. of two independent experiments, each acquiring three fields of view. *, p < 0.05; **, p < 0.01; ***, p < 0.001; ns, not significant.

The marked discrepancy between the level of IFITM3 upon combined CsA and CHX assessed by Western blotting (Fig. 1C) and by an indirect immunofluorescence assay (Fig. 1A, B) in digitonin-permeabilized cells implies that antibodies against the N-terminal segment of IFITM3 fail to detect the vast majority of IFITM3 molecules. Digitonin forms cholesterol-dependent pores in membranes and, thus, less efficiently permeabilizes cholesterol-poor membranes [33]. Cell permeabilization using a harsher permeabilizing agent, Triton X-100 (TX-100), which is largely independent of the lipid composition (reviewed in [33,34]), revealed a robust IFITM3 signal apparently associated with endosomes (Fig. 1D, E and see below). This signal is not considerably affected by CsA or CsA/CHX treatment. Such change in subcellular distribution is specific to IFITMs, since CsA/CHX treatment does not cause notable changes in the distribution or abundance of the Golgi markers, GM130 (Fig. 1A, B, D and E), Rab6, or TGN46 (Fig. S1).

The observed discrepancy in IFITM localization and abundance under different permeabilization conditions are not caused by IFITM3 overexpression in A549 cells. Similar effects of CsA treatment on the subcellular distributions of IFITM3 were observed in digitonin and TX-100 permeabilized HeLa cells endogenously expressing IFITM3 (Fig. S2).

To delineate the impact of membrane-permeabilization protocols on the apparent subcellular distribution of IFITM3 in CsA-treated cells, we used different permeabilizing agents (Figs. S3-S4). Streptolysin O, melittin, Tween20, and organic solvents (acetone and methanol) revealed various degrees of Golgi-associated IFITM3 signal in the presence of CsA (Figs. S3-S5). On the other hand, NP-40 permeabilization resulted in IFITM3 distribution that resembled that of TX-100. Thus, the apparent subcellular distribution of IFITM3 in CsA-treated cells is dependent on the harshness of membrane permeabilization.

We next assessed the impact of CsA treatment on the colocalization of IFITM3 with the late endosomes, where this protein normally accumulates [1,35,36]. A549.IFITM3 cells were permeabilized with digitonin or TX-100 and immunostained for IFITM3 and the marker for late endosomes, CD63 [37–39]. Whereas IFITM3 and CD63 poorly colocalized in digitonin-permeabilized cells treated with DMSO or CsA (Fig. 1G, I), these proteins colocalized well in TX-100 permeabilized cells exhibiting IFITM3 puncta distributed throughout the cells (Fig. 1H, I). We have observed a modest, but significant increase in colocalization of these proteins in CsA-treated samples, regardless of the permeabilization protocol. We also analyzed individual Z-stacks to minimize fortuitous colocalization due to signal overcrowding in maximum intensity projections (Fig. S6A-C).

To further verify that antibody access to the N-terminus of IFITM3 after CsA treatment is achieved through TX-100 treatment, but not digitonin permeabilization, we employed a two-step permeabilization and immunostaining protocol illustrated in Fig. S7A.First, A549.IFITM3 cells were permeabilized with digitonin, and accessible IFITM3 epitopes were saturated with rabbit anti-IFITM3 antibody followed by staining with secondary goat anti-rabbit antibodies. Next, cells were treated with TX-100 and incubated with excess of the same primary anti-IFITM3 antibody, followed by incubation with a differently labeled secondary goat anti-rabbit antibody. This protocol revealed two largely overlapping IFITM3 pools in DMSO-treated cells (Fig. S7B). However, cells pretreated with CsA contained two distinct IFITM3 pools accessible to antibodies through digitonin and TX-100 permeabilization. Whereas the IFITM3 signal after digitonin permeabilization was mainly concentrated in the perinuclear area, the additional IFITM3 signal appearing after TX-100 treatment was more peripherally distributed (Fig. S7B). After analysis of selected individual Z-stack images, we observed a change in colocalization of IFITM3 pools accessible by respective permeabilization step. The colocalization was higher in CsA-treated samples (Fig. S7C). In stark contrast, this 2-step immunofluorescence staining protocol did not reveal separate pools of CD63 (Fig. S7D, E). Parallel experiments using mouse anti-IFITM3 antibodies confirmed the existence of two IFITM3 pools with different antibody accessibility in CsA-treated cells (Fig. S7F, G). Collectively, our data support the existence of distinct pools of IFITM3 protein in CsA treated cells, differing in their accessibility to antibodies targeting the N-terminus; however, these pools provide no insight into the functional significance or underlying cause of this variation.

### CsA treatment occludes the IFITM’s N-terminal region

The masking of the N-terminus of IFITM3 in mildly permeabilized cells co-treated with CsA/CHX (Fig. 1) prompted us to test the accessibility of the C-terminus under these conditions. The absence of antibodies to a short C-terminal extracellular segment of IFITMs necessitates tagging of this protein. Since the C-terminus of IFITM3 is exposed to a degradative environment of late endosomes and lysosomes, C-terminally appended tags tend to be digested by proteases [4,5,26,35,40]. We, therefore, tagged the C-terminus of the plasma membrane-localized IFITM1 that faces the extracellular milieu [41–43]. A549 cells ectopically expressing IFITM1 fused with FLAG-tag at its C-terminus (A549.IFITM1-FLAG) were treated with CsA and permeabilized with digitonin or TX-100. Samples were co-immunostained using anti-IFITM1 (N-terminus) and anti-FLAG (C-terminus) antibodies, as illustrated in Fig. 2A. In control (DMSO-treated) cells permeabilized with digitonin or TX-100, the IFITM1’s N- and C-terminal signals largely colocalized at the plasma membrane, as expected (Fig. 2B, C). Strikingly, colocalization of the N- and C-terminal IFITM1 signals was significantly reduced in CsA-treated cells permeabilized with digitonin (Fig. 2B). The N-terminal signal concentrated in the perinuclear/Golgi area (as previously observed [32]), while the C-terminal signal appeared punctate, consistent with endosomal localization (Fig. 2B). By contrast, CsA-treated cells permeabilized with TX-100 exhibited good colocalization of N- and C-terminal signals that presumably localized to endosomes (Fig. 2C, D). We also analyzed individual Z-stacks to minimize fortuitous colocalization of abundant IFITM and CD3 signals in maximum intensity projection images (Fig. S6D). This analysis confirmed our initial observation of lower colocalization of N- and C-termini in digitonin permeabilized cells after treatment with CsA-compared to DMSO-treated cells; a higher colocalization was observed in TX-100 permeabilized cells.

**Figure 2.**
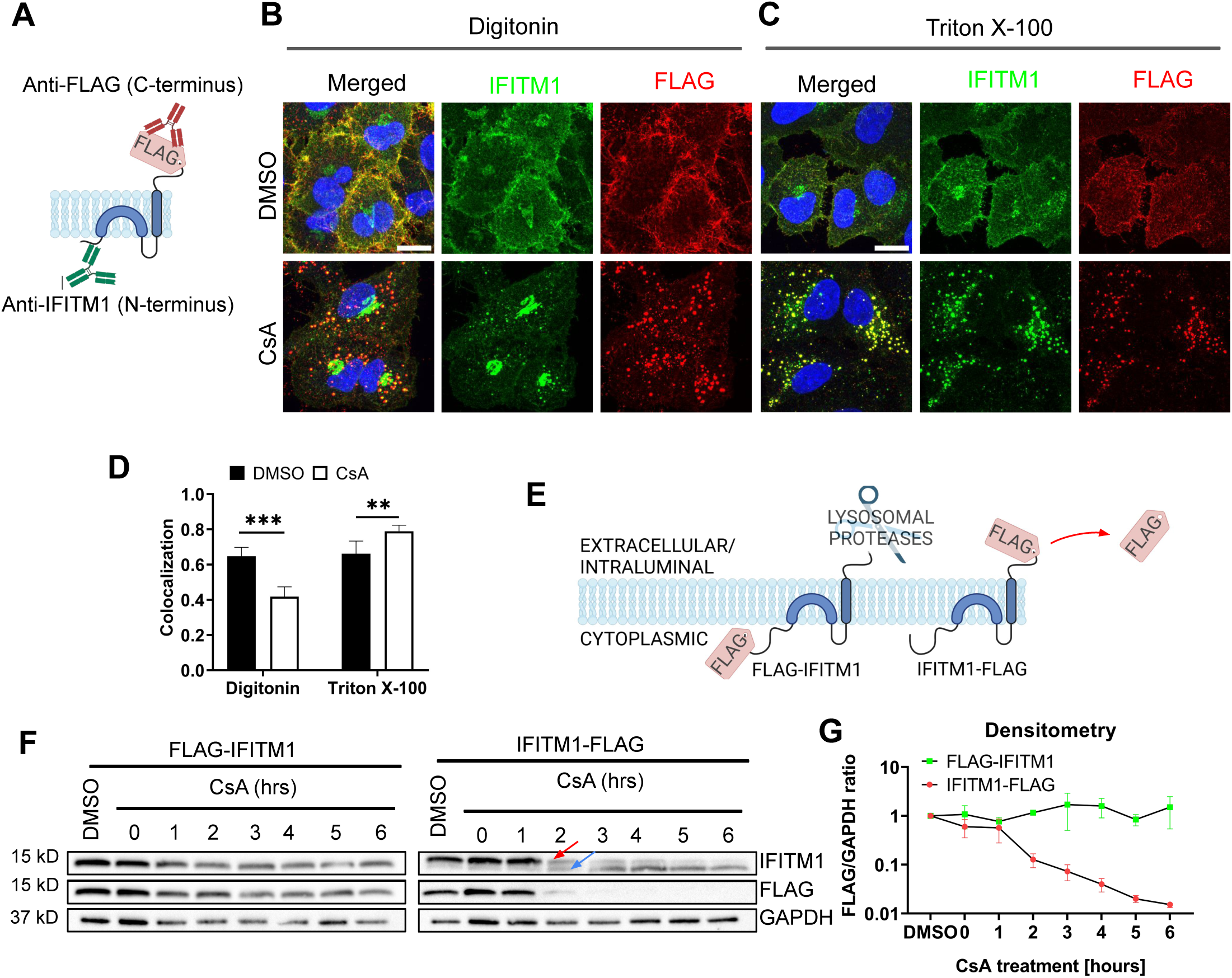
Disparate immunostaining patterns of the IFITM1’s N- and C-termini following CsA-treatment. (A) Illustration of a dual immunostaining strategy of IFITM1 fused to FLAG-tag at its C-terminus (IFITM1-FLAG). Not drawn to scale. (B, C) A549.IFITM1-FLAG cells were treated with DMSO or CsA (20 µM) for 90 minutes, fixed, permeabilized with either digitonin (B) or TX-100 (C), and stained using anti-IFITM1 (N-terminus, intracellular) or anti-FLAG (C-terminus, extracellular) antibodies. Scale bar 10 µm. (D) Colocalization of the IFITM1 N-terminus and C-terminal FLAG signals calculated using Mander’s overlap coefficient (MOC). Data are from two independent experiments. **, p < 0.01; ***, p < 0.001. (E) Illustration of IFITM1 protein with the FLAG-tag appended to the N-terminus (FLAG-IFITM1) or to the C-terminus (IFITM1-FLAG) and anticipated FLAG tag proteolysis in endolysosomes. (F) A549 cells ectopically expressing FLAG-IFITM1 or IFITM1-FLAG proteins were treated with DMSO or CsA (20 µM) for indicated times, harvested, and the cellular levels of IFITM1 and FLAG were assessed by Western blotting. (G) densitometry analysis of FLAG signal abundance (normalized to loading control, GAPDH). Red arrow points to the IFITM1-FLAG band, blue arrow points to the untagged IFITM1 band.

To test if IFITM1 relocalizes to late endosomes in the presence of CsA, A549.IFITM1-FLAG cells were pretreated with DMSO or CsA, permeabilized with digitonin or TX-100, and co-stained for CD63 and either IFITM1 N-terminus (using anti-IFITM1 antibody, Fig. S8A-C) or C-terminus (using anti-FLAG antibody, Fig. S8D-F). In CsA-treated and digitonin-permeabilized cells, the N-terminal signal largely concentrated in the perinuclear area, while the N- and C-terminal IFITM1 signals colocalized well with CD63 in TX-100 permeabilized CsA-treated cells (Fig. S8B, E). The IFITM1 C-terminus remains accessible to antibodies in digitonin-permeabilized cells. These observations led us to conclude that IFITM proteins are transported to late endosomes, where the N-terminus becomes poorly accessible to antibodies in mildly permeabilized cells through a yet unknown mechanism.

### CsA treatment does not change the IFITM’s membrane topology

Poor accessibility of the IFITMs’ N-terminal segment in CsA treated cells might be caused by changes in the protein’s structure and/or topology. It is generally accepted that IFITMs are single-span type II transmembrane proteins, with the N-terminus facing the cytosol and the C-terminus exposed to the extracellular milieu (IFITM1) or the lumen of endosomes (IFITM-2 and -3) [41,44]. Although this model is generally accepted, some studies suggested alternative topologies, including the inverted topology, with the N-terminus of IFITM proteins facing the extracellular space or lumen of endosomes [45,46].

To test possible CsA effects on IFITM1’s topology, we examined proteolysis of the N- and C-terminal FLAG-tags by Western blotting. This approach takes advantage of the IFITM1’s C-terminal tag cleavage by endosomal proteases after CsA-induced redistribution from the plasma membrane to late endosomes (Fig. 2E). We reasoned that a flipped topology would lead to clipping of the N-terminal FLAG tag by endosomal proteases. Cell lysates were analyzed by SDS-PAGE and blotted using anti-IFITM1 and anti-FLAG antibodies to distinguish protein degradation from selective FLAG cleavage. In both cell lines expressing N- and C-terminal FLAG-tagged IFITM1, a modest degradation of the IFITM1 protein was detected after a prolonged CsA treatment (Fig. 2F). However, only IFITM1-FLAG exhibited loss of FLAG signal in CsA-treated cells after 1 hour of treatment, with complete loss of FLAG signal after 3 hours. Loss of the FLAG tag was manifested by a concomitant increase in the IFITM1 band’s mobility (Fig. 2F, arrows), as expected. Importantly, we did not detect loss of the N-terminal FLAG tag at any point after CsA treatment (Fig. 2G).

To further probe possible changes in IFITM’s topology, we incubated A549.IFITM1-FLAG cells with CsA overnight and chased in a CsA-free growth medium which lacked or contained CHX to block protein synthesis, as shown in Fig. S9A. After incubation for up to 6 hours, samples were harvested and examined by Western blotting. We observed a slow recovery of the FLAG signal with a concurrent shift of an untagged IFITM1 band to a FLAG-tagged IFITM1 band starting at 3 hours after CsA removal (Fig. S9B). As expected, the FLAG signal recovery was blocked in the presence of CHX. Together, these results argue against possible CsA-induced changes in IFITM’s topology.

To verify that clipping of C-terminal FLAG occurs in endolysosomes, we co-treated cells with CsA and either the lysosomal pathway inhibitors, Bafilomycin A1 (BafA1) and NH_4_Cl, or proteasomal degradation inhibitors, MG132 and Lactacystin. Cells were also co-treated with a pan-cathepsin inhibitor, E64-d. Co-treatment with CsA/BafA1 or CsA/NH_4_Cl abrogated the IFITM1’s mobility shift and concomitant loss of FLAG signal (Fig. S9C). By comparison, partial inhibition was observed in cells co-treated with non-specific proteasome inhibitor MG132, while co-treatment with a more specific inhibitor Lactacystin did not inhibit FLAG removal from IFITM1. Inhibition of lysosomal cathepsins by E64-d showed only partial inhibition on CsA-driven FLAG loss (Fig. S9C). The activity of the MG132 and Lactacystin was confirmed by blotting using an anti-ubiquitin antibody. As expected, both MG132 and Lactacystin induced the accumulation of ubiquitinated proteins due to the block of the proteasomal pathway (Fig. S9C).

The above results show that the C-terminus of IFITM1 in CsA-treated cells is facing the lumen of late endosomes, implying that the topology of this protein is not altered compared to cells’ basal condition.

### CsA rescues IAV fusion with IFITM-expressing cells through a mechanism that is distinct from those of rapamycin and MK-2206

As reported previously by us and others [29,30,32], rapamycin antagonizes the IFITM3’s antiviral activity. Shi et al. concluded that rapamycin leads to IFITM3 degradation through inhibition of mTOR and subsequent phosphorylation of TFEB, the master regulator of lysosome function and microautophagy. However, this effect seems to require the N-terminus of IFITM3, since rapamycin fails to promote degradation of the Δ17-20 IFITM3 mutant, which lacks YEML endocytic motif, localizes to the plasma membrane, and restricts a different set of viruses [30]. Indeed, the IAVpp fusion block was relieved by rapamycin in A549.IFITM3 cells but only partially recovered in A549.IFITM1-FLAG cells (Fig. 3A).

**Figure 3.**
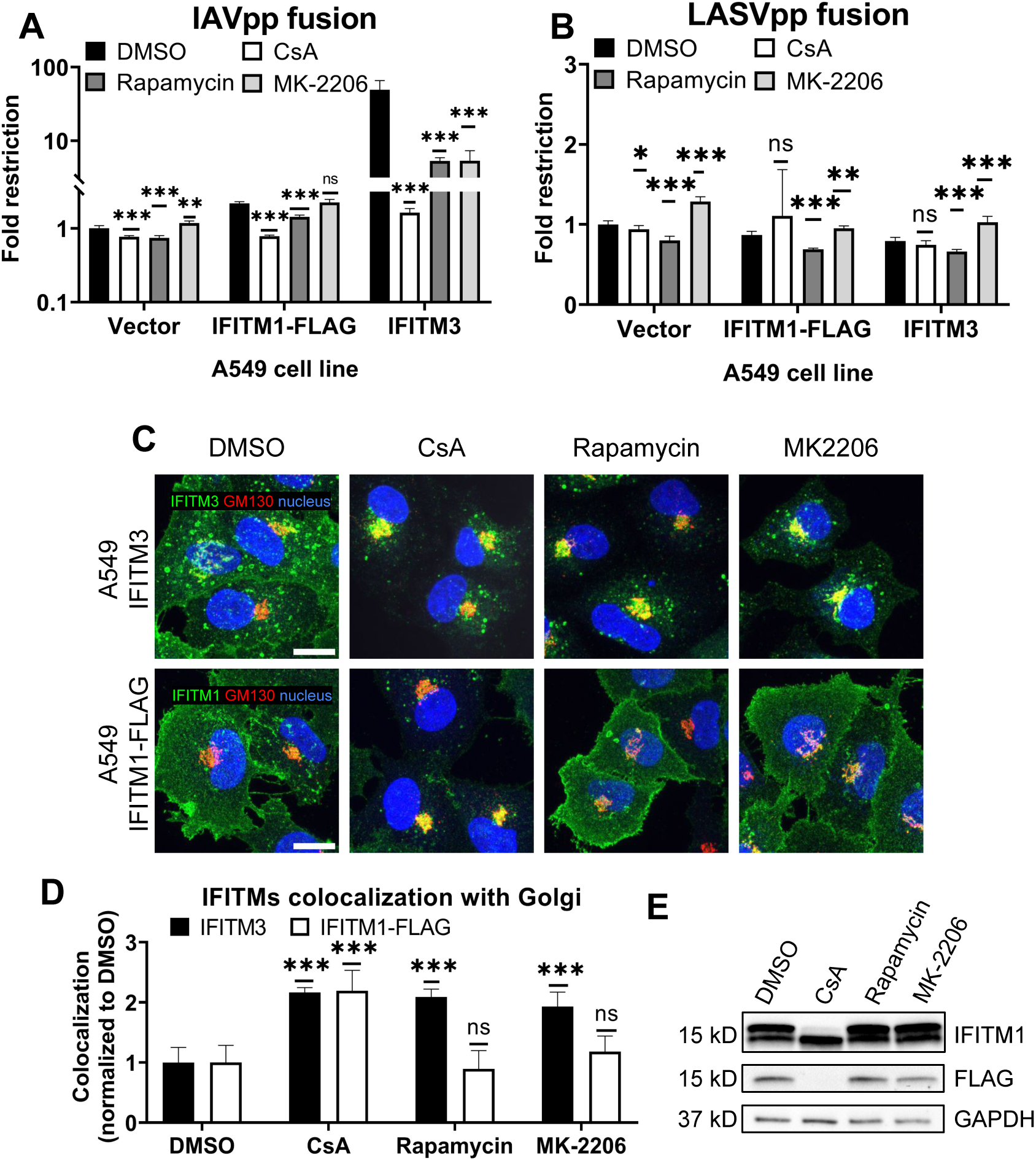
CsA-induced rescue of IAV fusion with IFITM1 expressing cells occurs through a mechanism that is distinct from those of rapamycin and MK-2206. (A, B) A549.Vector, A549.IFITM3 or A549.IFITM1-FLAG cells were preincubated in the presence of DMSO, CsA (20 µM), rapamycin (20 µM), or MK-2206 (10 µM) for 90 minutes and challenged with IAVpp (A) or LASVpp (B) pseudoviruses, and viral fusion was measured using a beta-lactamase assay. (C) A549.IFITM3 or A549.IFITM1-FLAG cells were treated as in (A), fixed, stained for GM130 and respective IFITM proteins, and imaged. Scale bar 10 µm. (D) Colocalization between IFITMs and GM130 in cells shown in (C) was calculated using MOC. (E) A549.IFITM1-FLAG cells were treated as in (A), harvested, and cell lysates were analyzed by Western blotting. Data are means and S.D. of two independent experiments, each performed in triplicate. *, p < 0.05; **, p < 0.01; ns, not significant.

During our screening of inhibitors of various cellular pathways that might antagonize IFITM3, we found that the Akt inhibitor, MK-2206, rescued IAV-cell fusion in A549.IFITM3 cells (Fig. 3A). Note that both rapamycin and MK-2206 had non-specifically modulated fusion of LASV pseudoviruses, which are resistant to IFITM-mediated restriction [1] (Fig. 3B). This effect may be due to inhibition of the PI3K/AKT/mTOR pathway. We also observed a modest, yet statistically significant, drop in viability in cells treated with the above compounds (Fig. S10). Interestingly, unlike CsA, neither rapamycin nor MK-2206 induced relocalization of IFITM1-FLAG protein from the plasma membrane, while both successfully altered the subcellular distribution of IFITM3 (Fig. 3C, E). Finally, only CsA treatment of A549.IFITM1-FLAG cells caused loss of FLAG and concomitant shift in IFITM1 band mobility on immunoblots (Fig. 3E).

Taken together, our data suggest a fundamentally different mechanism of CsA action on IFITMs that, in contrast to rapamycin and MK-2206, modulates the subcellular distribution of both IFITM3 and IFITM1 and potently enhances virus-cell fusion.

### CsA treatment sequesters IFITMs inside late endosomes, likely within intraluminal vesicles

Our results (Figs. 1 and 2) reveal that CsA treatment relocalizes IFITM1 to late endosomes, while, except for the newly synthesized pool of IFITM3, this protein remain largely endosome-associated. In both cases, CsA treatment leads to selective masking of the protein’s N-terminus in cells permeabilized with digitonin, without a change in IFITM’s membrane topology. This surprising observation can be explained by IFITM1 and IFITM3 redistribution from the PM and the limiting membrane (LM) of late endosomes, respectively, to intraluminal vesicles (ILVs) of multivesicular bodies (MVBs), which are complex and dynamic structures (reviewed in [47,48]). To test the notion that the inaccessibility of ILVs to digitonin is the reason for poor immunostaining of IFITMs in CsA-treated cells, we employed the epidermal growth factor receptor (EGFR) as a reference marker. EGFR is a type I transmembrane protein that is redirected from the plasma membrane to ILVs upon activation by the EGF ligand [49,50]. We took advantage of the ability to immunolabel the extracellular and intracellular domains of EGFR and IFITM1-FLAG independently to examine the accessibility of respective epitopes in digitonin permeabilized cells (Fig. 4A). A549.IFITM1-FLAG cells were treated, as shown in Fig. 4B. Briefly, cells were pretreated with CHX for 1 hour to block protein synthesis, exposed to either EGF or CsA on ice for 30 min, and shifted to 37 °C. Samples were fixed at indicated times, permeabilized with digitonin, and stained for extracellular domains (N-terminus of EGFR and C-terminus of IFITM1-FLAG) and intracellular domains (C-terminus of EGFR and N-terminus of IFITM1-FLAG).The weak and dispersed signal of EGFR is likely due to inhibition of the requisite EGFR dimerization in the cold [51,52]. The EGFR aggregation and internalization from the plasma membrane occurred within 10 minutes, while IFITM1 internalization was detectable at ∼20 minutes after shifting to 37 °C (Fig. 4C, D). Both proteins showed a marked shift from the plasma membrane to endosomal compartments, along with strongly diminished signals of their respective intracellular domains after 60 minutes of treatment with EGF or CsA (Fig. 4E-G). These data suggest that both proteins are redistributed to the ILVs upon CsA treatment, as the signal of their extracellular domains weakened over time when compared to the respective signal from intracellular domains. These results support our model that, in CsA-treated cells, the extracellular domains are facing the lumen of MVBs, which is accessible to antibodies in digitonin-permeabilized cells, while the intracellular domains are hidden inside the ILVs.

**Figure 4.**
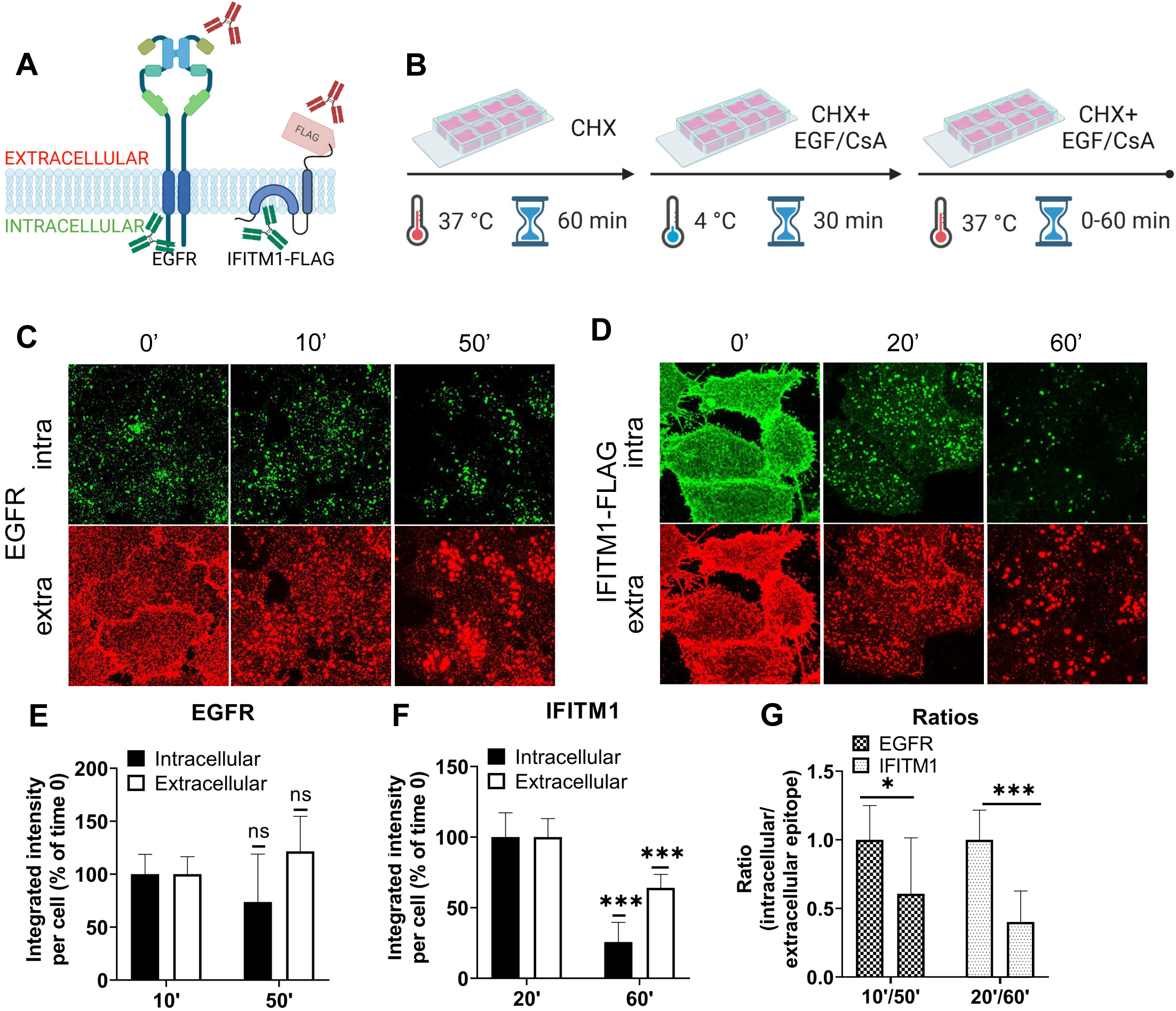
The IFITMs’ N-terminal region is selectively sequestered in late endosomes of CsA-treated cells. (A) Antibodies recognizing extracellular or intracellular domains of EGFR or IFITM1-FLAG proteins were used to probe the accessibility of these domains in cells treated with the EGFR ligand or CsA. Not drawn to scale. (B) A549 cells were incubated in the presence of CHX for one hour, placed on ice, treated with EGF (EGFR samples) or CsA (IFITM1 samples) for 30 minutes, and returned to 37 °C for indicated times. Cells were fixed, permeabilized with digitonin, and stained for extracellular and intracellular domains of a respective target protein, EGFR (C) or IFITM1/FLAG (D). The integrated intensity for each respective antibody targeting domains of EGFR (E) or IFITM1 (F) was calculated and plotted as a function of time of incubation. (G) The ratios between integrated intensities of intracellular and extracellular domains of EGFR and IFITM1 at indicated times are plotted. Data are means and S.D. of two independent experiments, each acquiring three fields of view. ***, p < 0.001; ns, not significant.

We visualized the dynamics of IFITM1 internalization in the presence of CsA by directly labeling IFITM1-C-FLAG with anti-FLAG antibody conjugated to AlexaFluor 647 and performed live cell imaging. IFITM1 was rapidly relocalized from the plasma membrane (Movie S2), which was not observed in control experiments (Movie S1). The aggregation of IFITM1 signal in cytosolic puncta started around 10 min time and culminated at 20 min.

To investigate the mechanism of CsA-induced IFITM1 internalization, inhibitors targeting macropinocytosis (EIPA [53,54]) and dynamin-dependent endocytosis (Dynasore [55]) were employed. As expected, EIPA and Dynasore inhibited the uptake of respective cargoes – 70 kDa dextran (macropinocytosis) and transferrin (clathrin-mediated endocytosis) (Fig. S11A). Notably, Dynasore had minimal impact on CsA-induced IFITM1 internalization, whereas EIPA significantly disrupted IFITM1 relocalization to late endosomes (Fig. S11B-D). It should be pointed out that EIPA did not fully block CsA-induced internalization of IFITM1, as this protein’s colocalization with the plasma membrane stained with WGA was significantly reduced in EIPA/CsA samples compared to EIPA/DMSO samples (Fig. S11B). We note that co-treatment with CsA and these inhibitors—especially EIPA—mildly reduced cell viability. Interestingly, our markers for macropinocytosis (Dextran) and clathrin-mediated endocytosis (EGF) showed high colocalization after 30 min of CsA treatment, suggesting that both pathways eventually converge, which is in line with published studies (reviewed in [56]).

To further test if IFITM1 is relocalized to ILVs by CsA, we employed a super-resolution stimulated emission depletion (STED) microscopy of IFITMs and EGFR, a well-established ILV marker upon ligand (EGF) binding (reviewed in [57]). A549.IFITM1-C-FLAG cells were pretreated with CHX for 1 hour prior to incubation with a combination of CHX, EGF, and CsA (or DMSO as control) on ice for 30 minutes (similar to Fig. 4B). Cells were then shifted to 37 °C for 1 hour, fixed, permeabilized with TX-100, and stained using anti-IFITM1 antibodies targeting the intracellular epitope N-terminal region of IFITM1, and anti-EGFR antibodies, targeting the extracellular epitope. While there was no colocalization between IFITM1 and EGFR signals in mock treated cells (Fig. S12A), these proteins appeared to colocalize in CsA-treated cells (Fig. S12B), suggesting a convergence of these proteins in the same pool of endosomes.

Lastly, we assessed whether CsA induces IFITM3 relocalization from the LM to ILVs using STED microscopy. A549.IFITM3 (IFITM3+) or control A549.vector (IFITM3-) cells were treated using the protocol described above for IFITM1-C-FLAG STED experiments and in Fig. 4B. Cells were fixed, permeabilized with TX-100, and stained using anti-IFITM3 antibody targeting the intracellular epitope, and anti-EGFR, targeting the extracellular epitope. In control A549.IFITM3 cells treated with DMSO, endosomes tended to have a hollow, doughnut-shaped appearance based upon the peripherally localized EGFR signal, with a diameter of 1.2±0.2 µm; IFITM3 and EGFR partially colocalized at the periphery of these endosomes (Fig. 5A). Notably, most of the EGFR signal was punctate. In contrast, CsA treatment reduced the diameter of endosomes to 0.68±0.17 µm, and these endosomes were “filled” with the IFITM3 that was no longer localized to the LM (Fig. 5B). We did not observe enlarged endosomes or the effect of CsA on their size in IFITM3-negative control cells (Fig. 5C, D) regardless of the treatment (0.7±0.2 µm vs 0.6±0.2 µm for DMSO and CsA treated cells, respectively).

**Figure 5.**
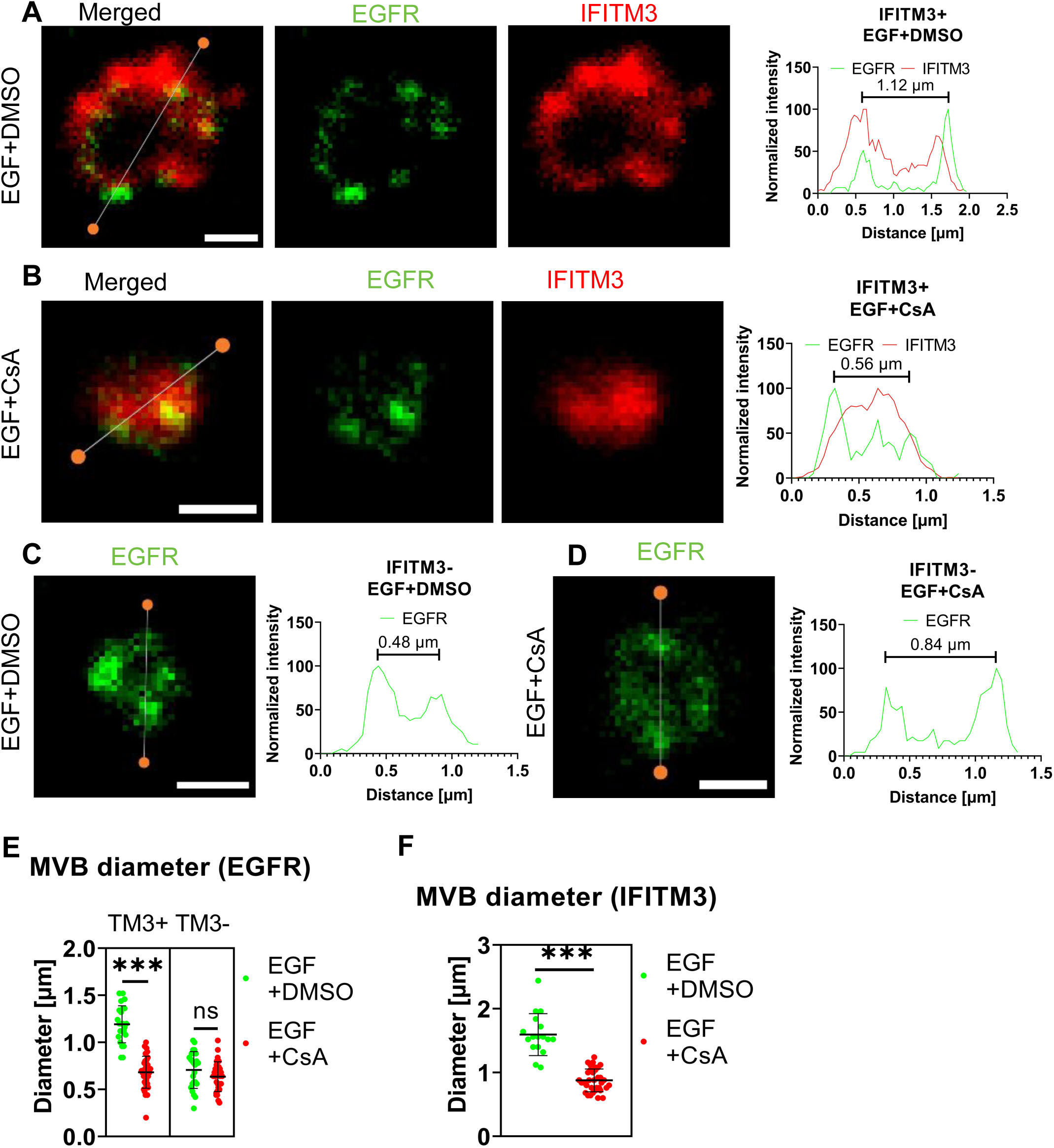
IFITM3 relocates to the interior of late endosomes upon CsA treatment. A549.IFITM3 (IFITM3+) or A549.vector (IFITM3-) cells were incubated in the presence of CHX for one hour, placed on ice, treated with EGF and either DMSO (A, C) or CsA (B, D) for 30 minutes, and returned to 37 °C for indicated times. Cells were fixed, permeabilized with TX-100, and stained using anti-EGFR (targeting N-terminus) and anti-IFITM3 primary antibodies and secondary antibodies conjugated to STED-compatible fluorophores, STAR RED and STAR 580. Normalized linear intensity profiles across endosomes are shown for each channel. To measure the endosome diameter, local maxima of EGFR signals were used. Endosomes with low EGFR signal, excessive background noise, or indistinguishable features were excluded. Representative linear histograms for IFITM3 positive or negative cells in the presence or absence of CsA are shown. (E) Endosome diameters based on EGFR signal are plotted. Endosomes from two independent experiments (n>20 endosomes, n>15 cells) were analyzed per condition and per cell line (A549.IFITM3 and A549.vector). (F) The endosome diameters based on IFITM3’s signal for IFITM3+ cells were calculated based upon the distance between the normalized linear profile intensities corresponding to 25% of signal. Endosomes with a high background were omitted. Lines and bars are medians and interquartile range. Scale bar is 0.5 µm. ***, p < 0.001; ns, not significant.

**Figure 6.**
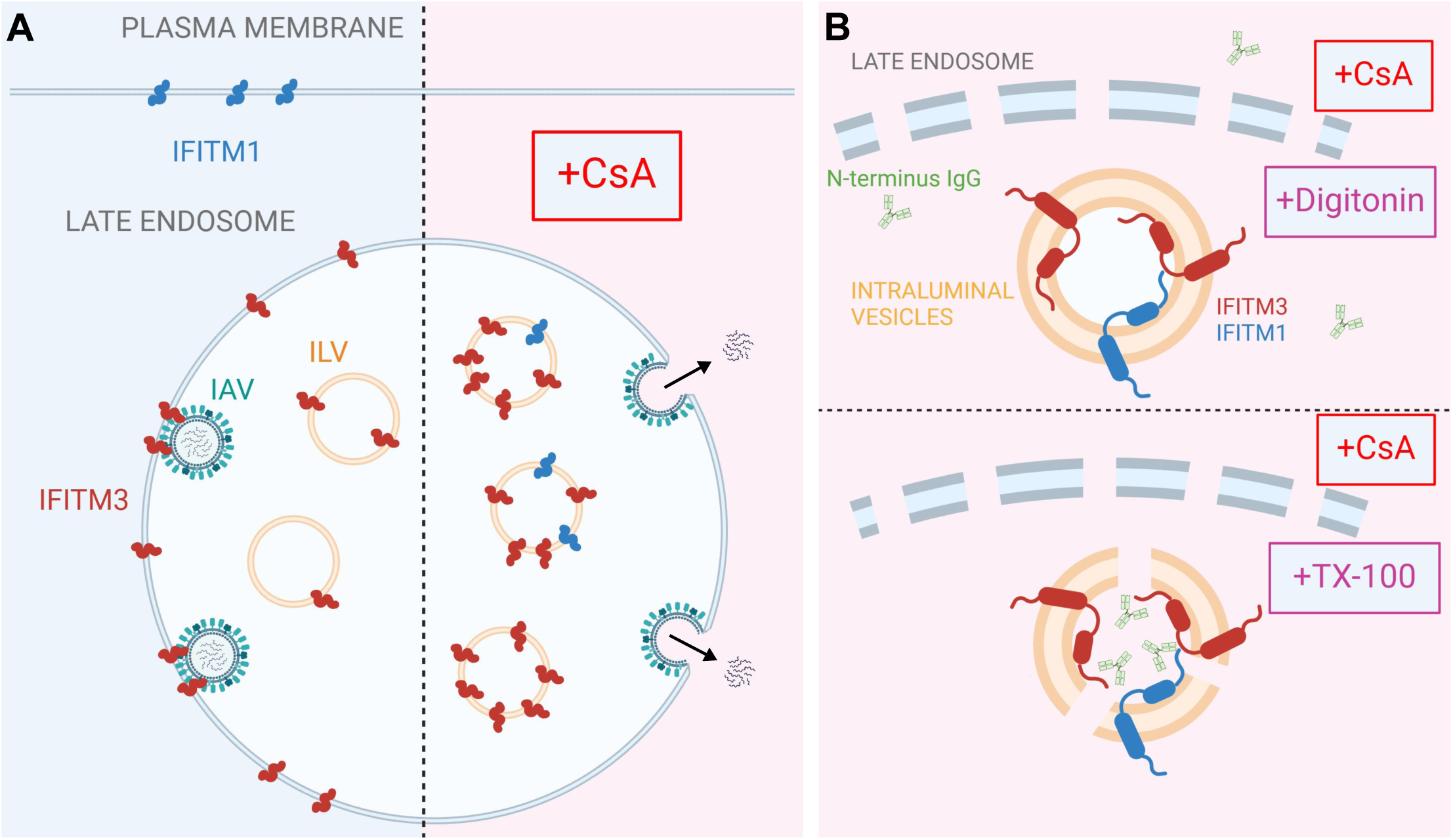
A model for CsA-induced IFITM relocalization to the ILVs of late endosomes and rescue of IAV fusion. A proposed model of modulation of subcellular localization of IFITMs by CsA. (A) In the absence of CsA, IFITM1 is primarily located at the plasma membrane, while IFITM3 concentrates in the limiting membrane and intraluminal vesicles (ILVs) of the late endosome. This basal subcellular distribution of IFITM proteins inhibits the fusion of incoming viruses (e.g. IAV) with the cell membranes. CsA redirects IFITM proteins from their respective locations to the ILVs, sequestering them away from incoming viruses and allowing viral fusion to occur at the respective cellular locations. (B) Digitonin does not permeabilize ILVs, thus precluding access of antibodies to the N-terminus of IFITM proteins. In contrast, TX-100 disrupts the ILV membrane, enabling antibody binding to the N-terminus of IFITM proteins. Visual representations are not drawn to scale. Created in BioRender.com.

Alternatively, we infected A549 cells with AF-568-labeled IAV in the presence or absence of CsA. To achieve non-invasive labeling, we used IFITM3-iSNAP in combination with SNAP-Cell 647-SiR [58]. IAV was found to colocalize with the limiting membrane of late endosomes, as marked by the IFITM3-iSNAP signal (Fig. S13A). CsA-induced changes in the distribution of IFITM3-iSNAP and IAV that clearly shifted from the limiting membrane toward the center of the endosome (Fig. S13B).

## Discussion

While IFITMs play an important role in curbing viral infection, the mechanism of their antiviral activity is not fully understood. Hurdles to delineating the mechanism of IFITM action include uncertainty regarding their membrane topology and complex regulation of their subcellular localization by single residue substitutions and post-translational modifications [23–28]. Pleiotropic effects of CsA on cellular processes [59–61] precluded the identification of factors/pathways regulating IFITMs’ localization and antiviral activity in treated cells. The results reported in this study provide new insights into the mechanism of CsA-mediated rescue of viral fusion through regulation of IFITMs’ trafficking/localization (Fig. 7).

Co-treatment with CsA and CHX (to eliminate the signal from newly synthesized IFITM pool that traffics through the Golgi) revealed a large pool of both IFITM1 and IFITM3 in endosomes. Importantly, this pool is only detectable by immunofluorescence in TX-100 permeabilized cells that gains access to ILVs, whereas mild permeabilization with digitonin allows antibody access almost exclusively to the Golgi-trapped pool of IFITMs. Indeed, there is evidence that overexpressed IFITM3 accumulates in the Golgi and delays transport of other glycoproteins transport through this apparatus [62]. The impact of CsA on IFITM1 localization is particularly striking, since unlike the endosome-localized IFITM3, IFITM1 is nearly fully relocalizes to late endosomes/ILVs.

Two lines of evidence support the notion that CsA induces IFITM1 and IFITM3 redistribution from the plasma membrane and the limiting membrane of late endosomes, respectively, to ILVs.

First, the N-terminus of IFITMs is selectively sequestered in intracellular compartments that are not accessible to antibodies in digitonin-permeabilized cells. This effect is similar the sequestration of the cytoplasmic tail of EGFR, a well-characterized protein targeted to ILVs upon ligand (EGF) binding (reviewed in [57]). Importantly, the lack of CsA’s effect on the overall level and topology of IFITMs in A549 cells rules out partial or full cleavage of the N-terminal region recognized by the antibodies as the reason for loss of the immunofluorescence (IF) signal. Second, super-resolution microscopy implies that IFITM3 is translocated from the LM of late endosomes to the lumen, and this relocalization is associated with shrinking of the endosome’s diameter. This finding is also supported by our observation that C-terminal FLAG-tag on IFITM1 is cleaved in the presence of CsA.

ILVs originate from the LM of MVBs and carry cargo destined for degradation, secretion, or temporary segregation from the cytoplasm (reviewed in [63]. Importantly, IFITM1 or IFITM3 proteins trapped in ILVs after CsA treatment are not degraded for hours, as evidenced by constant levels of these proteins in cell lysates (Fig. 1C). It is worth noting that endogenously expressed IFITM3 in HeLa cells treated with CsA appears to be degraded within a few hours of CsA treatment [32]. The ILV formation is regulated by the ESCRT machinery, with ALIX and TSG101 playing key roles [64–66]. These proteins can perform partially overlapping functions confounding the results of knockdown experiments. While the ESCRT system is central to ILV biogenesis, studies have shown ILV production in cells lacking multiple ESCRT proteins, indicating the contribution of endosomal lipids, BMP and ceramide, in ILV biogenesis [67–69]. It was also reported that IFITM3 expression affects cholesterol levels and distribution, either directly [70] or through inhibition of VAMP-Associated Proteins [71].

CsA is known to partition into and alter the properties of lipid membranes, including shifting the phase transition temperature and lipid domain morphology [72–74]. CsA also selectively interacts with sphingomyelin [75]. Given that the antiviral activity of IFITMs is modulated by their interactions with lipids, such as cholesterol and phosphoinositides [76–79], it is conceivable that CsA can also modulate the subcellular distribution of IFITMs indirectly, through modifying the cell membranes. However, this mechanism does not fully explain how IFITM1 is transported from the plasma membrane to the LM of late endosomes and then to ILVs, suggesting the involvement of additional host cofactors. It is intriguing that rapamycin and MK-2206, both inhibitors of the PI3K/AKT/mTOR (PAM) pathway, do not impact the localization of IFITM1 or the N-terminally truncated IFITM3 lacking the endocytic signal [30]. This suggests that CsA may influence multiple pathways that exert broader effects on IFITMs and cellular processes. However, the non-specific effects of PAM inhibitors, rapamycin and MK-2206, on cell viability could reduce virus-cell fusion, potentially leading to decreased fusion efficiency.

CsA-mediated redistribution of IFITMs has implications beyond viral entry and infection. IFITM proteins play a role in cancer, syncytiotrophoblast fusion, and inhibition of ILV back-fusion [80–84]. The rapid and non-toxic redistribution of IFITMs by CsA offers a promising means to counteract the above adverse effects of IFITMs and improve lentivirus-based gene delivery [31]. Unlike Rapamycin and MK-2206, CsA successfully redistributes IFITMs, which increases its utility for modulating adverse effects of these proteins. Future studies will be aimed at identifying the IFITM motif and cellular partners responsible for the rapid and selective translocation into ILVs. This knowledge can be utilized for a controlled sequestration of target cellular proteins into ILVS.

## Material and methods

### Cell Lines, Plasmids, and Reagents

Human A549, HEK293T/17, and HeLa cells were obtained from ATCC (Manassas, VA, USA) and grown in Dulbecco’s Modified Eagle Medium (DMEM) supplemented with 10% heat-inactivated fetal bovine serum (FBS, Atlanta Biologicals, Flowery Branch, GA, USA), 100 U penicillin-streptomycin (Gemini Bio-Products, Sacramento, CA, USA). Stable cell lines A549.vector, A549.IFITM1-FLAG, A549.FLAG-IFITM1, A549.IFITM3, and A549.IFITM3-iSNAP ectopically expressing the respective IFITM proteins have been described previously [58,85]. Briefly, cells were transduced with VSV-G-pseudotyped viruses encoding wild-type or flag-tagged IFITMs or with the empty vector, pQCXIP (Takara, Shiga, Japan), and selected with 1.5 µg/mL puromycin.

Bafilomycin A1 (Cat. B1793), NH_4_Cl (Cat. A0171), E64d (Cat. E8640), Cyclosporine A (Cat. 30024), cycloheximide (Cat. C7698), rapamycin (Cat. 553210), Triton X-100, Streptolysin O, melittin, and acetone were from Sigma (St. Louis, MO, USA). MG-132 (Cat. 474791) was purchased from Calbiochem (Columbus, OH, USA). Lactacystin (sc-3575) was purchased from Santa Cruz Biotech (Dallas, TX, USA). Recombinant human EGF (Cat. 236-EG) was purchased from R&D Systems (Minneapolis, MN, USA). Nonidet P-40 was purchased from USBiological (Salem, MA, USA), Tween 20 was obtained from J.T. Baker (Phillipsburg, NJ, USA), methanol was from Fisher Chemicals (Zurich, Switzerland), and Digitonin was purchased from Invitrogen (Cat. 11024-24-1, Research product international, Mount Prospect, Illinois). MK-2206 was from Selleckchem (Cat. S1078, Houston, TX). SNAP-Cell 647-SiR (Cat. S9102S) and SNAP-Cell Oregon Green (Cat. S9104S) were purchased from New England Biolabs (Ipswich, MA, USA). FM1-43 dye (Cat. T35356) and Alexa Fluo 568 NHS Ester (Cat. A20103) were purchased from Invitrogen (Waltham, MA, USA). mCLING labeled with ATTO 647N (Cat. 710 006AT647N) was obtained from Synaptic Systems (Goettingen, Germany). The Influenza A/PR/8/34 virus (Cat. 10100374) was purchased from Charles River Laboratories (Wilmington, MA, USA).

Antibodies used were rabbit IgG against the N-terminus of IFITM3 (Abgent, San Diego, CA, USA), mouse anti-IFITM2/3 (Proteintech, San Diego, CA, USA), rabbit anti-IFITM1 (Sigma), mouse anti-GM130 (BD Bioscience, Franklin Lakes, NJ, USA), sheep anti-TGN46 (Bio-Rad AbD Serotec Limited, Luxembourg), mouse anti-Rab6A, clone 5B10 (a gift from Prof. Angelika Barnekow, Münster University, Germany), mouse anti-flag® M2 (Sigma), mouse anti-Human CD63 (BD Biosciences), Influenza A NP recombinant rabbit monoclonal antibody (Fisher) and antibody to the EGFR N-terminus (Calbiochem), antibody to the EGFR C-terminus (Cell Signaling, Danvers, MA, USA), AlexaFluor 568 Goat anti-Mouse IgG (H+L) (Invitrogen, Waltham, MA, USA), Goat anti-rabbit IgG (H+L) conjugated with AlexaFluor 647 (ThermoFisher, Waltham, MA, USA), and Donkey anti-sheep IgG (H+L) conjugated with AlexaFluor 568 (Abcam). Secondary antibodies conjugated with STED-compatible dyes were STAR RED (STRED-1002) and STAR 580 (ST580-1001), both purchased from Abberior, Germany.

### Pseudovirus production

Pseudovirus production protocols and plasmid information were described previously [32]. Briefly, HEK297T/17 cells were transfected using JetPRIME transfection reagent (Polyplus-transfection, New York, NY). For Influenza A pseudoviruses (IAVpp), ∼70% confluent cells in a 100-mm tissue culture dish were transfected with 5 µg pR9deltaEnv, 1.5 µg pMM310 plasmid encoding Vpr fused to β-lactamase, 1 µg pcRev, and with envelope glycoprotein-encoding plasmids: pCAGGS WSN HA and NA (2.5 µg each) plasmids. For Lassa pseudoviruses (LASVpp), 4 μg Lassa-GPC encoding plasmid was used instead of HA and NA. After 12 hours, the transfection medium was replaced with a phenol red-free growth medium, and cells were cultured for 36 hours, at which point, the medium was collected, filtered through 0.45 µm PES membrane filter (VWR, Radnor, PA), concentrated 10× using Lenti-X™ Concentrator (Clontech, Mountain View, CA), and stored in aliquots at −80 °C.

### IAV labeling, purification, and characterization

Twenty-five µL of freshly prepared 1 M sodium bicarbonate (pH 9.0) buffer was mixed with 75 µL of ultrapure water to make the reaction solution. Fifty µL of IAV (2 mg/mL of total protein) was mixed with 100 µL of reaction solution and incubated for 1 hour at room temperature with AF568-NHS at a concentration of 50 µM by agitating in the dark at the lowest speed of a vortex. After incubation, NHS activity was quenched by adding 3 µL of 1 M Tris-buffer (pH 8.0). Unbound dye was removed using NAP-5 gel filtration column (Illustra, Danaher Corporation, USA) according to the manufacture’s manual. Labeled IAV was eluted with 500 µL of PBS without calcium and magnesium (PBS –/–; 21-040-CV, Corning), and filtered through a 0.45 µm filter. Labeled IAV was frozen and stored at −80 °C.

To assess the effect of IAV labeling on virus titer, 10^5^ A549 cells were seeded in each well of 96-well plate and cultured overnight. Next day, unlabeled IAV and IAV-AF568 stocks were serially diluted with DMEM supplemented with 2% FBS (DMEM/2% FBS) and spinoculated onto A549 cells at 4 °C, 1500xg for 30 minutes. Cells were washed with fresh medium to remove unbound viruses and cultured in DMEM/2%FBS at 37 °C for ∼20 hours. Cells were then fixed with 4% PFA (ThermoFisher) for 15 min at room temperature, permeabilized with 0.3% Triton X-100 for 15 min, blocked with 10% FBS for 1 hour, and incubated with 10 μg/mL of Influenza A NP antibody at room temperature for 2 hours, followed by labeling with 2 μg/mL of Goat anti-Rabbit IgG–FITC antibody at room temperature for 45 min. Cell nuclei were labeled with 10 µM of Hoechst 33342 at room temperature for 10 min. Immunostained cells were imaged with BioTek Cytation 5 Cell Imaging Multimode Reader (BioTek Instruments, Agilent Technologies, USA). The infected cells were counted to determine the viral titer.

### Western Blotting

Cells were harvested and processed, as described elsewhere [86]. Proteins were detected with rabbit anti-IFITM3, rabbit anti-IFITM1, mouse anti-FLAG, mouse anti-Ubiquitin (P4D1, Santa Cruz), or mouse anti-GAPDH (Proteintech) antibodies and horseradish peroxidase-conjugated Protein G (VWR), using a chemiluminescence reagent from Cytiva (Marlborough, MA, USA). The chemiluminescence signal was detected using an XR+ gel doc (Bio-Rad, Hercules, CA, USA). Densitometry was performed using Image lab (version 3.0, Bio-Rad).

### BlaM assay

The β-lactamase (BlaM) assay for virus–cell fusion was carried out, in a modified version of a previously described method [86]. Briefly, cells were pretreated with respective drug at given concentration for 90 minutes, after which pseudovirus bearing respective envelope glycoprotein and β-lactamase fused to Vpr (BlaM-Vpr) was bound to target cells plated in black clear-bottom 96-well plates by centrifugation at 4 °C for 30 min at 1550× g. Unbound viruses were removed by washing, and fusion was initiated by shifting cells to 37 °C, 5% CO_2_ for 120 min, after which time cells were loaded with the CCF4-AM BlaM substrate (Life Technologies). The cytoplasmic BlaM activity (ratio of blue to green fluorescence) was measured after overnight incubation at 12 °C, using a Synergy HT fluorescence microplate reader (Agilent Bio-Tek, Santa Clara, CA, USA). Cell viability was determined using the CellTiter-Blue Reagent (Promega); after adding this reagent to cells, the samples were incubated for 30 to 60 min at 37 °C, 5% CO_2_, and cell viability was measured on Synergy HT plate reader (579_Ex_/584_Em_).

### Endocytosis Inhibition by Pharmacological Drugs

For dextran uptake assay, A549.IFITM1-C-FLAG cells were preincubated with DMSO, EIPA (50 μM) or Dynasore (120 μM) for 30 min. We added 150 μg/mL tetramethylrhodamine dextran (TMR-dextran, ThermoFisher Scientific, D1818, 70,000 MW) to cells and incubated at 37 °C for 30 min. Dynasore treated cells were kept in serum-less medium.

For transferrin uptake measurements, A549.IFITM1-C-FLAG cells were pretreated with DMSO, EIPA or Dynasore. Dynasore treated cells were kept in serum-less medium. Cells were kept on ice for 5 min, and Transferrin-fluorescein (Transferrin from Human Serum, Fluorescein Conjugate, ThermoFisher Scientific, T35352, 50 μg/mL) was added and incubated on ice for 15 min. Unbound transferrin was removed by two PBS washes, and the cells were placed at 37 °C for 10 min. EIPA or Dynasore were maintained in medium throughout the experiment (during preincubation, washing, and post-incubation). Cells were transferred to ice, chilled for 5 minutes, washed with PBS and fixed with 4% paraformaldehyde. Samples were blocked using 10% FBS for 30 minutes and stained with anti-Flag antibody conjugated with AF-647.

For CsA co-treatment, cells were preincubated in fresh medium for 45 minutes. After that, cells were transferred on ice and allowed to cool down for 5 minutes prior the 30 minutes pharmacological drug and EGF treatment and anti-Flag antibody conjugated with AF-647, after which the medium was changed for DMSO- or CsA-containing medium and cells were shifted to 37C for 30 minutes. After this, cells were washed with PBS (containing respective drug) and fixed with 4% paraformaldehyde. Samples were blocked using 10% FBS for 30 minutes and stained with Wheat Germ Agglutinin (WGA) Alexa Fluor 568 Conjugate (Biotiuim, 29077-1) to label the cell membrane. Fluorescence intensity was measured using a 561 nm laser line for Dextran-TMR or transferrin-fluorescein AF-555, and a 633 nm laser line for WGA imaging.

### Immunostaining, microscopy, live cell imaging, and image analyses

One day before imaging, cells were plated in 8-well chamber coverslips (Nunc, Rochester, NY, USA) coated with 0.2 mg/mL collagen (Cat. C9791, Sigma). Cells were treated with indicated compounds/inhibitors or left untreated, fixed with 4% PFA (ThermoFisher) for 20 min at room temperature, permeabilized with 150 µg/mL digitonin or 0.1% Triton X-100 for 20 min, and blocked with 10% FBS for 30 min. Cells were next incubated with respective primary antibodies diluted in 10% FBS for 1.5 h, washed, and incubated with secondary antibodies in 10% FBS for 45 min. Samples were stained with Hoechst 33342 (4 µM, Invitrogen) in PBS for 5–10 min before imaging.

Cells used for consecutive permeabilization by digitonin and Triton X-100 were treated with DMSO or CsA (20 µM) for 90 minutes, fixed with 4% PFA for 20 min at room temperature, permeabilized with 150 µg/mL digitonin, and blocked with 10% FBS for 30 min. Cells were next incubated with anti-IFITM3 (Abgent) antibodies diluted in 10% FBS for 1.5 h, washed, and incubated with anti-rabbit secondary antibodies conjugated with AF647 in 10% FBS for 45 min. Next, cells were permeabilized with 0.1% TX-100 for 20 min and blocked with 10% FBS for 30 min. Cells were next incubated with anti-IFITM3 (Abgent) antibodies diluted in 10% FBS for 1.5 h, washed, and incubated with anti-rabbit secondary antibodies conjugated with AF568 in 10% FBS for 45 min. Cell nuclei were stained with Hoechst 33342 (4 µM, Invitrogen) in PBS for 5–10 min before imaging.

For live cell imaging, cells were seeded on collagen-coated glass-bottom dishes (MatTek, Ashland, MA) day before the experiment in phenol red-less medium. The next day, cells were chilled on ice and incubated with anti-Flag antibody conjugated with AF-647 for 30 minutes. After that, cells were incubated in the presence of Hoechst 33342 (4 µM, Invitrogen) for 10 min before imaging at room temperature, washed with pre-warmed Live Cell Imaging Solution (LCIS, Invitrogen) twice. Cells in 1 ml of LCIS were moved to a pre-warmed microscope chamber and allowed to equilibrate at 37 °C before 1 ml of LCIS containing either DMSO or 50 µM of CsA was added. The acquired time-lapse (acquisition every 10 second) Z-stack (10) images were converted to maximum intensity projections.

Images were acquired on a Zeiss LSM 880 confocal microscope using a plan-apochromat 63×/1.4NA oil objective. The entire cell volume was imaged by collecting multiple Z-stacks. Images were analyzed using FIJI [87]. Protein signal colocalization (using both Pearson’s and Mander’s coefficients) was computed by the JaCoP FIJI plugin [88] on maximum-intensity projection images. For 3D analysis, individual Z-stacks capturing the bottom half of the cells were analyzed using the JaCop FIJI plugin.

### CsA/EGF CHX chase protocol

Cells were incubated in the presence or absence of 10 μg/mL CHX for 1 hour at 37 °C, placed on ice, and treated with combinations of CHX with CsA (20 µM) or EGF (50 ng/mL) in HEPES-buffered medium for 30 minutes on ice. Cells were shifted to 37 °C for various times before fixation with 4% PFA, permeabilization with digitonin, and immunostaining.

### STED imaging and analysis

Cells were incubated in the presence of 10 μg/mL CHX for 1 hour at 37 °C, placed on ice, and treated with HEPES-buffered medium containing combinations of CHX and CsA (20 µM) or CHX and EGF (50 ng/mL) for 30 minutes. Cells were shifted to 37 °C for 1 hour before fixation with PFA, permeabilization with 0.1% TX-100, and subsequent staining using respective primary followed by secondary antibodies conjugated to STED-compatible fluorophores.

In IFITM3-iSNAP and IAV imaging experiments, A549.IFITM3-iSNAP cells were pre-incubated with DMSO or 20 µM of CsA for 1.5 hours and spinoculated with AF-568 labeled IAV at MOI of 2 at 4 °C, 1500x g for 30 minutes. Infection was allowed to proceed for 1 hour in the presence of DMSO or CsA, at which time, cells were stained with SNAP-Cell 647-SiR for 30 min, washed and incubated with fresh medium for additional 30 min to remove unbound dye. Cells were fixed with 4% PFA for STED super-resolution microscopy.

STED Facility Line super-resolution microscope (Abberior) on an inverted Olympus IX83 body using 60×/1.42NA oil objective, two excitation laser lines (561 nm and 640 nm), and two pulsed STED lasers (595 nm and 775 nm, respectively) were used for imaging. The entire volume of selected endosomes was imaged by collecting multiple Z-stacks at 50 nm intervals, with a pixel size of 50 nm. Line histograms across endosomes were drawn, and histograms of normalized intensity were used to assess IFITM3 or IFITM1 and EGFR distribution within endosomes. Endosomal IFITM3-iSNAP and IAV particles were segmented in 3D by MorphoLibJ Fiji plugin, and the distance between individual IAV and the center of the endosome was measured by 3D manager Fiji plugin and normalized to the endosome’s radius.

### Statistical Analysis

Unpaired Student’s t-test or Mann-Whitney test using GraphPad Prism version 9.3.1 for Windows (GraphPad Software, La Jolla, CA, USA), as indicated.

## Acknowledgments

The authors would like to thank the members of Melikian lab members, Mariana Marin, Gokul Raghunath, Smita Verma, and Sergii Buth, for reading the manuscript and valuable discussions. We also wish to thank Hui Wu and Monica Macias for technical support. We are grateful to Dr. Baek Kim for access to BioTek Cytation 5. Research reported in this publication was supported by the NIAID R01 grant AI135806 to G.B.M.

## Supplemental Figure

**Figure S1.**
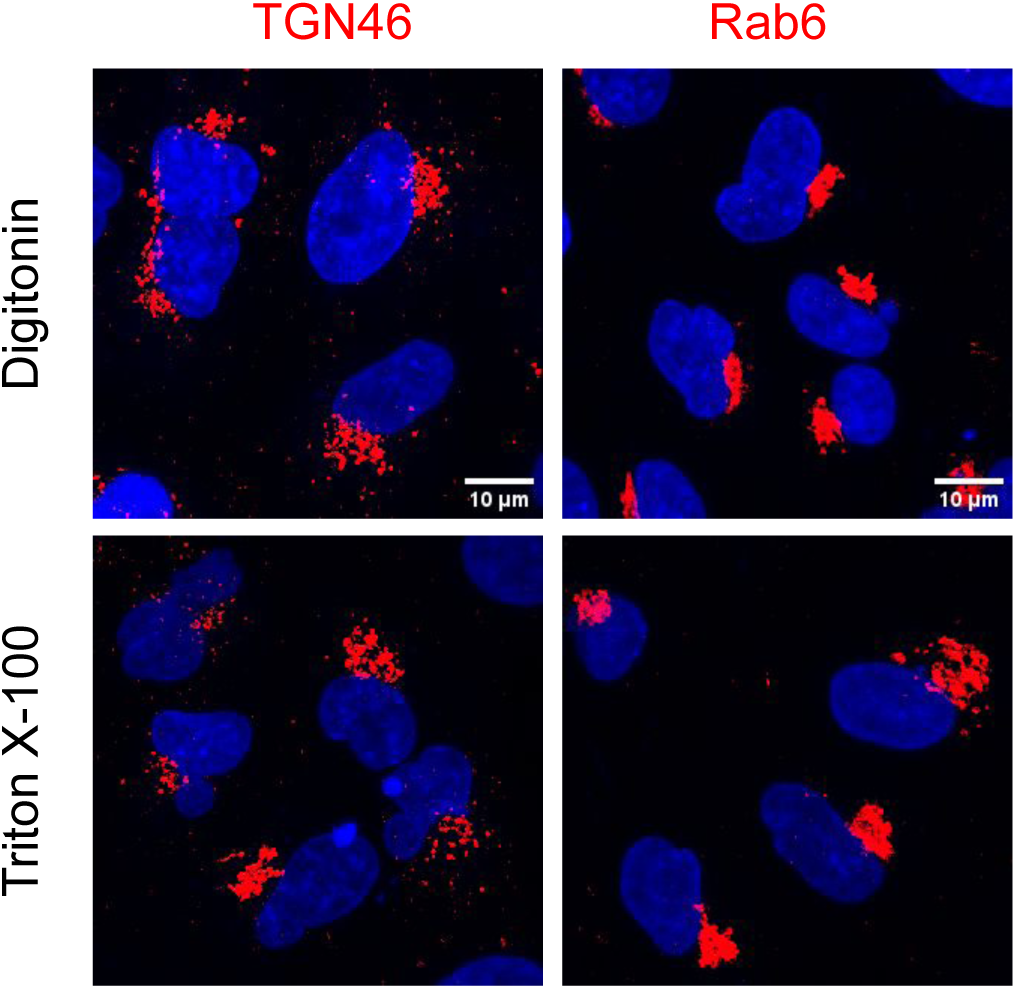
TGN46 and Rab6 subcellular localization in cells permeabilized by digitonin or Triton X-100. A549.IFITM3 cells were fixed, permeabilized with either digitonin or TX-100, and stained with either anti-TGN46 or anti-Rab6 antibodies.

**Figure S2.**
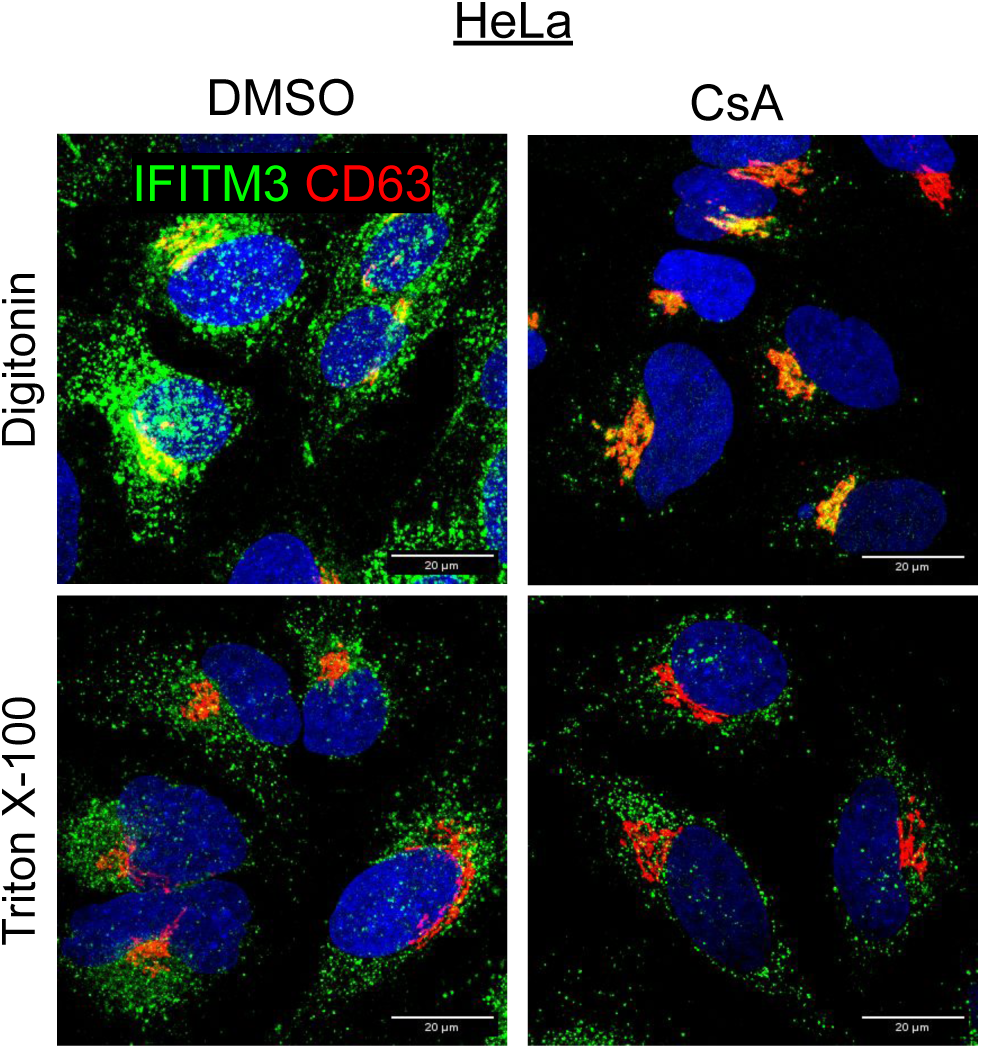
Different permeabilization protocols of HeLa cells revealed two pools of IFITM3. HeLa cells were incubated in the presence or absence of CsA (20 µM) for 90 minutes, fixed, permeabilized with either digitonin or TX-100 and stained with anti-IFITM3 and anti-GM130 antibodies.

**Figure S3.**
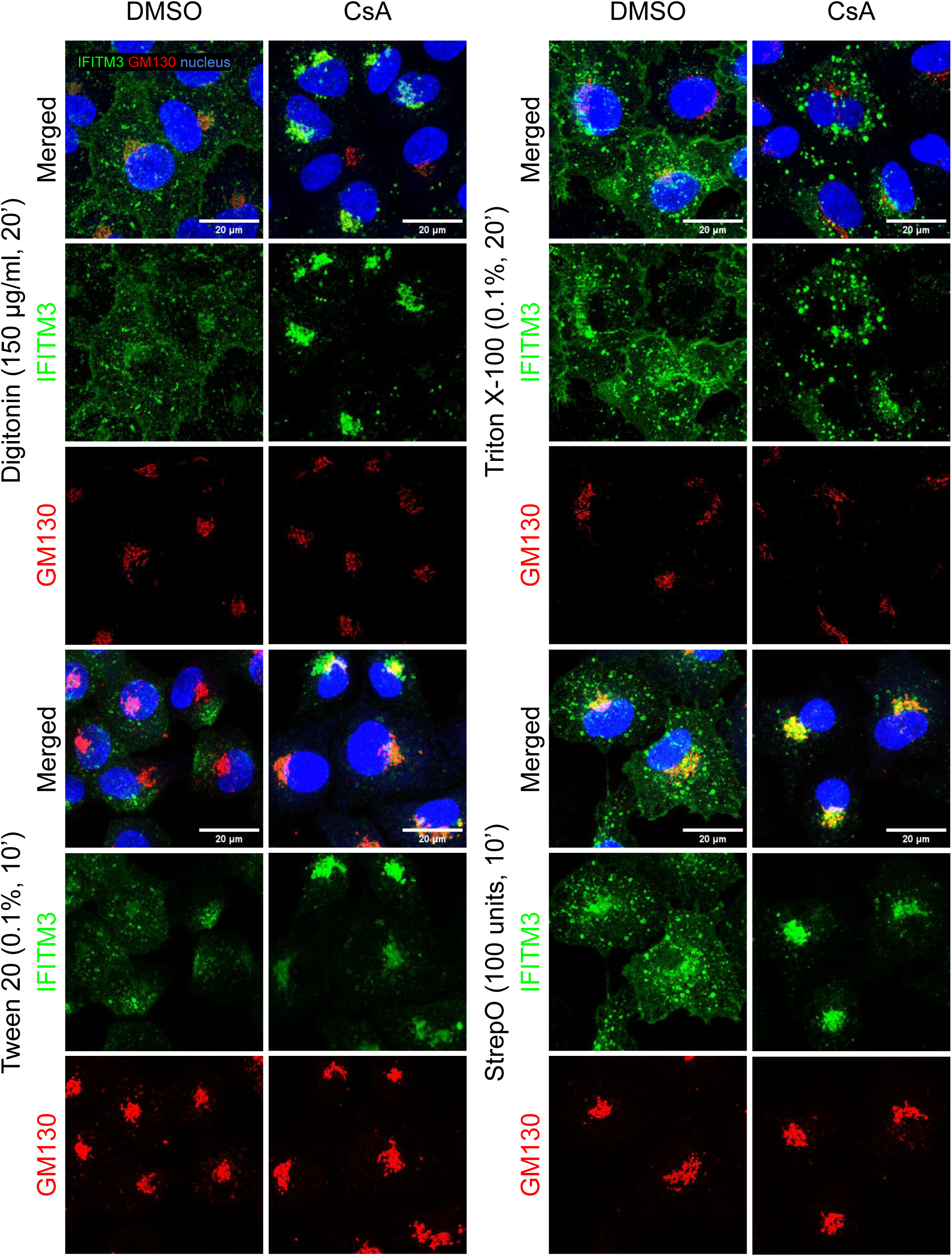
Subcellular distribution of IFITM3 in control and CsA treated cells following different A549.IFITM3 cell permeabilization protocols. A549.IFITM3 cells were incubated in the presence or absence of CsA (20 µM) for 90 minutes, fixed with PFA, and permeabilized with different reagents, as indicated, followed with staining using anti-IFITM3 and anti-GM130 antibodies.

**Figure S4.**
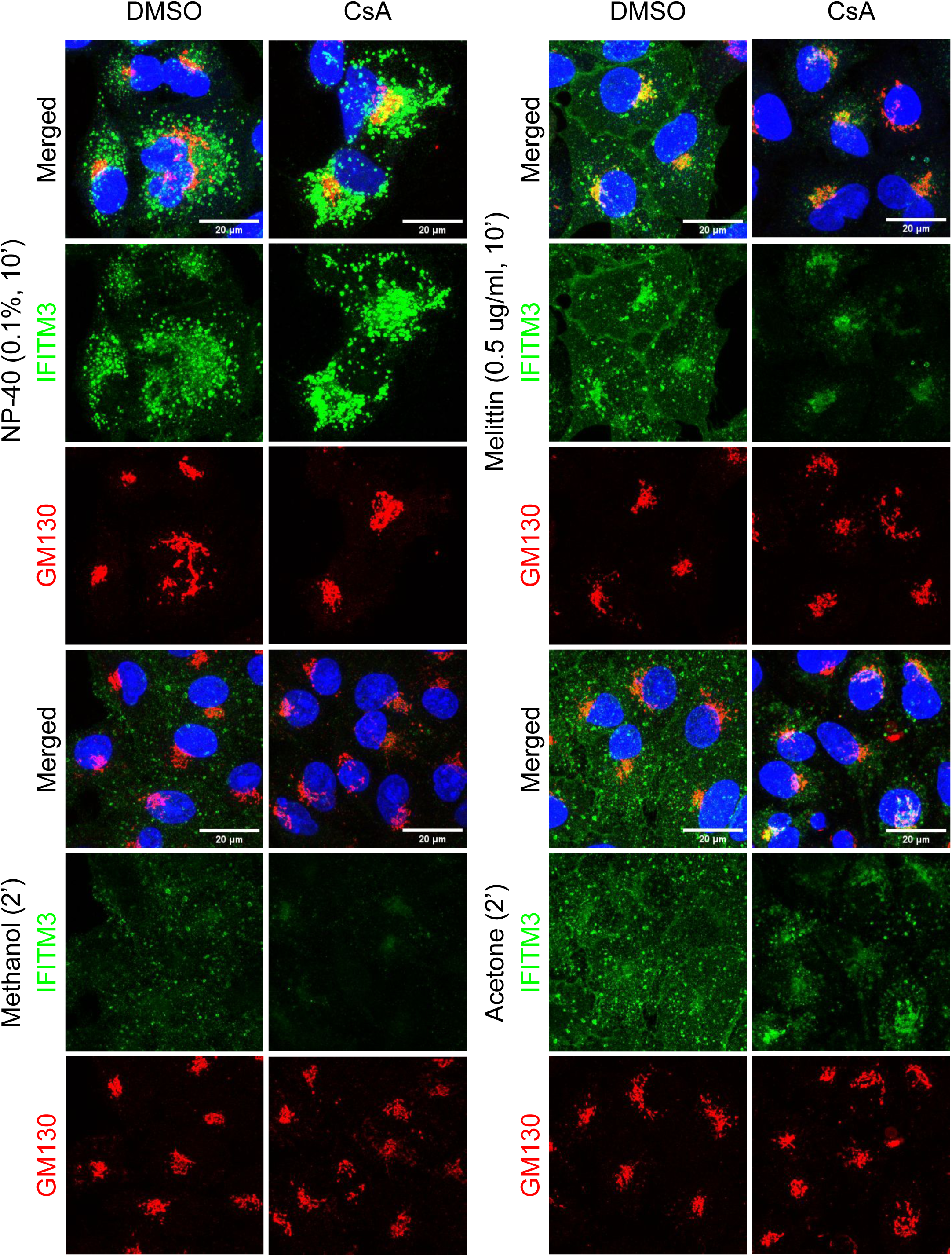
Subcellular distribution of IFITM3 in control and CsA treated cells following different A549.IFITM3 cell permeabilization protocols. A549.IFITM3 cells were incubated in the presence or absence of CsA (20 µM) for 90 minutes, fixed with PFA, and permeabilized with different reagents, as indicated, followed with staining using anti-IFITM3 and anti-GM130 antibodies.

**Figures S5.**
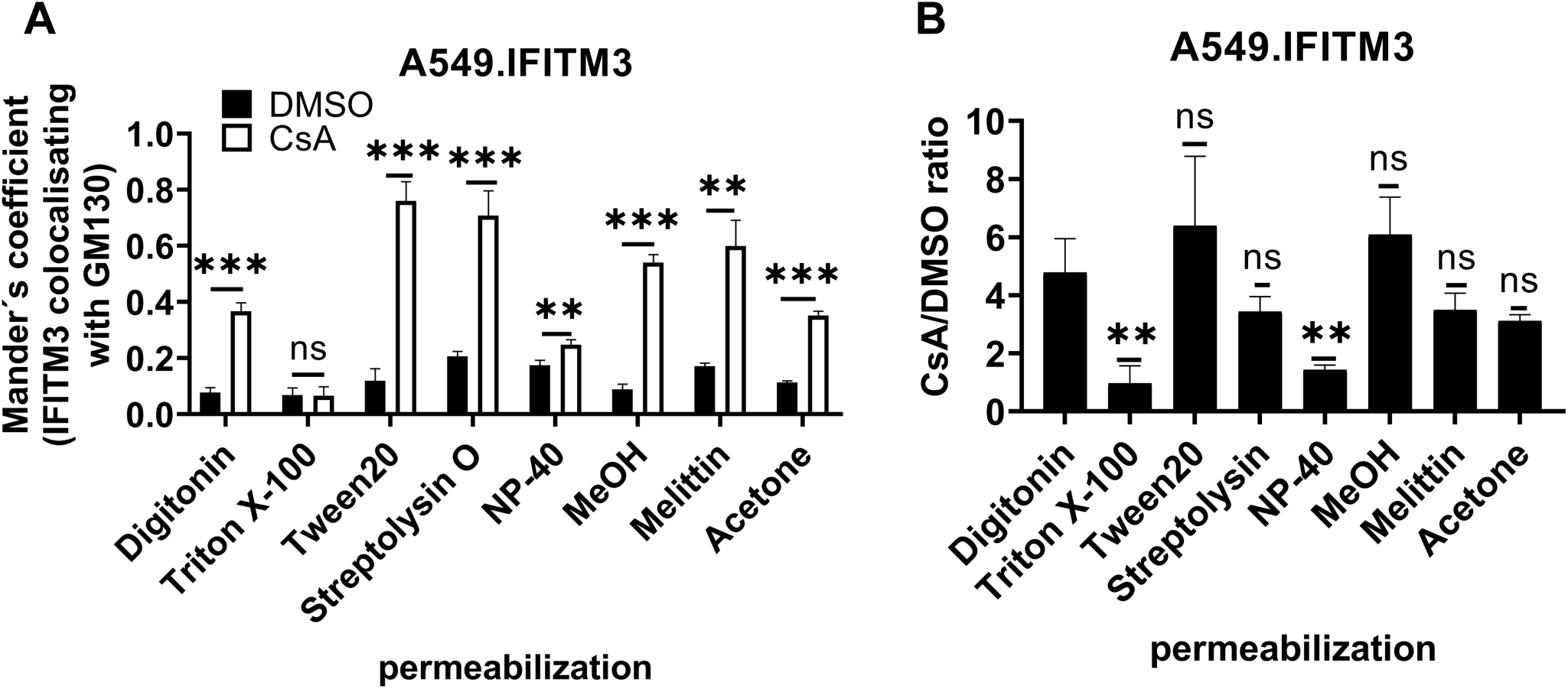
Analysis of IFITM3 and GM130 colocalization in A549.IFITM3 cells permeabilized with different reagents. (A) Colocalization (Mander’s coefficient) of IFITM3 and GM130 for cells treated with DMSO or CsA, as well as the ratio between colocalization in DMSO and CsA-treated (see Figs. S4 and S5) was calculated using the JaCoP FiJi plugin. (B) Ratios of IFITM3/GM130 colocalization in CsA vs DMSO treated cells calculated from the results in panel (A). Statistical significance of ratios between Digitonin sample and respective sample was obtained by computing the z-score. ** p < 0.01; *** p < 0.001; ns, not significant.

**Figure S6.**
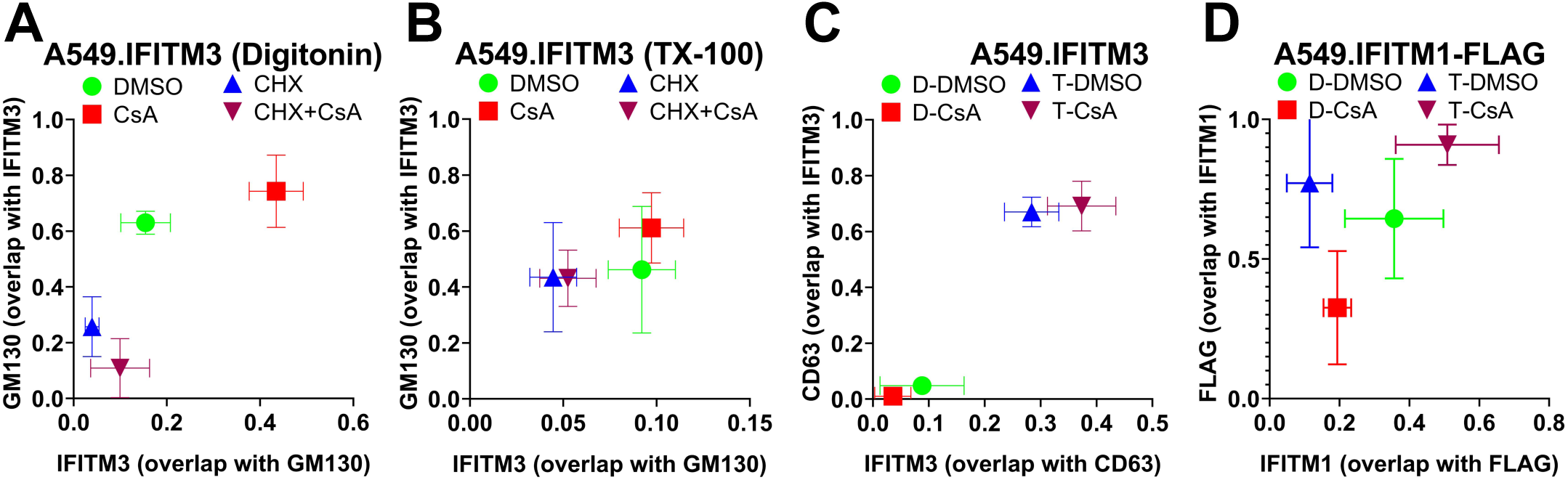
3D analysis of IFITM1 and CD63 signal colocalization. Colocalization analysis of selected individual Z-stacks representing the lower half of A549.IFITM3 (A-C) or A549.IFITM1-FLAG (D) cells. Reciprocal Mander’s overlap coefficients are plotted on the axis. This analysis relates to the main figures Fig. 1A, digitonin permeabilization) or Fig. 1D (TX-100 permeabilization) showing colocalization between IFITM3 and CD63 (Fig. 1G, H), and between IFITM1 and FLAG (Fig. 2C). Colocalization was determined slice-by-slice, means and error bars are the collection of data from respective slides. Data are means and S.D. of two independent experiments, each containing three fields of view. Measured Mander’s Overlap Coefficients (MOC) were plotted with IFITM3 signal overlap with GM130 or CD63 on the X-axis and GM130/CD63 overlap with IFITM3 on the Y-axis.

**Figure S7.**
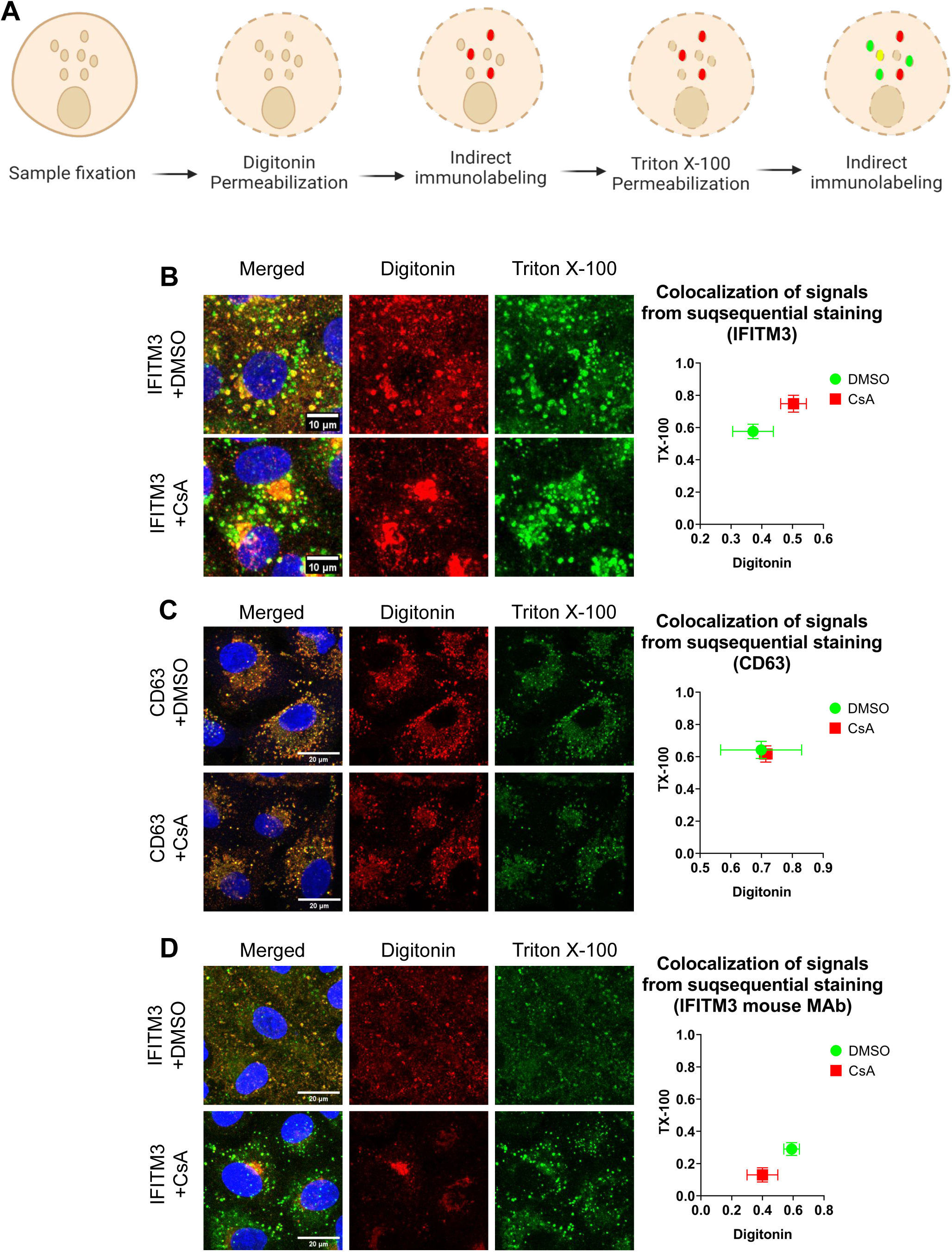
Distinct pools of IFITM3 revealed by different permeabilization protocols. (A) Illustration of consecutive immunostaining steps following cells permeabilization with digitonin and TX-100. (B) A549.IFITM3 cells were treated with DMSO or CsA, fixed, permeabilized with digitonin, incubated with rabbit anti-IFITM3 antibody, then permeabilized with TX-100, and incubated with mouse anti-IFITM3 antibody. Primary antibody binding was detected using different secondary antibodies conjugated to different fluorophores, to distinguish IFITM3 proteins recognized in the respective permeabilization steps. (C) A549.IFITM3 cells were treated as in (B), but an anti-CD63 antibody was used to visualize the CD63 pools accessible after each permeabilization step. (D) A549.IFITM3 cells were treated as in (B), but a mouse anti-IFITM3 antibody was used to detect IFITM3. All colocalizations of respective signals were determined by MOC on individual slices slice-by-slice (analyzed as in Fig. S6). Digitonin signals overlapping with TX-100 signals were plotted on the X axis, TX-100 signals overlapping with digitonin were plotted on the Y axis.

**Figure S8.**
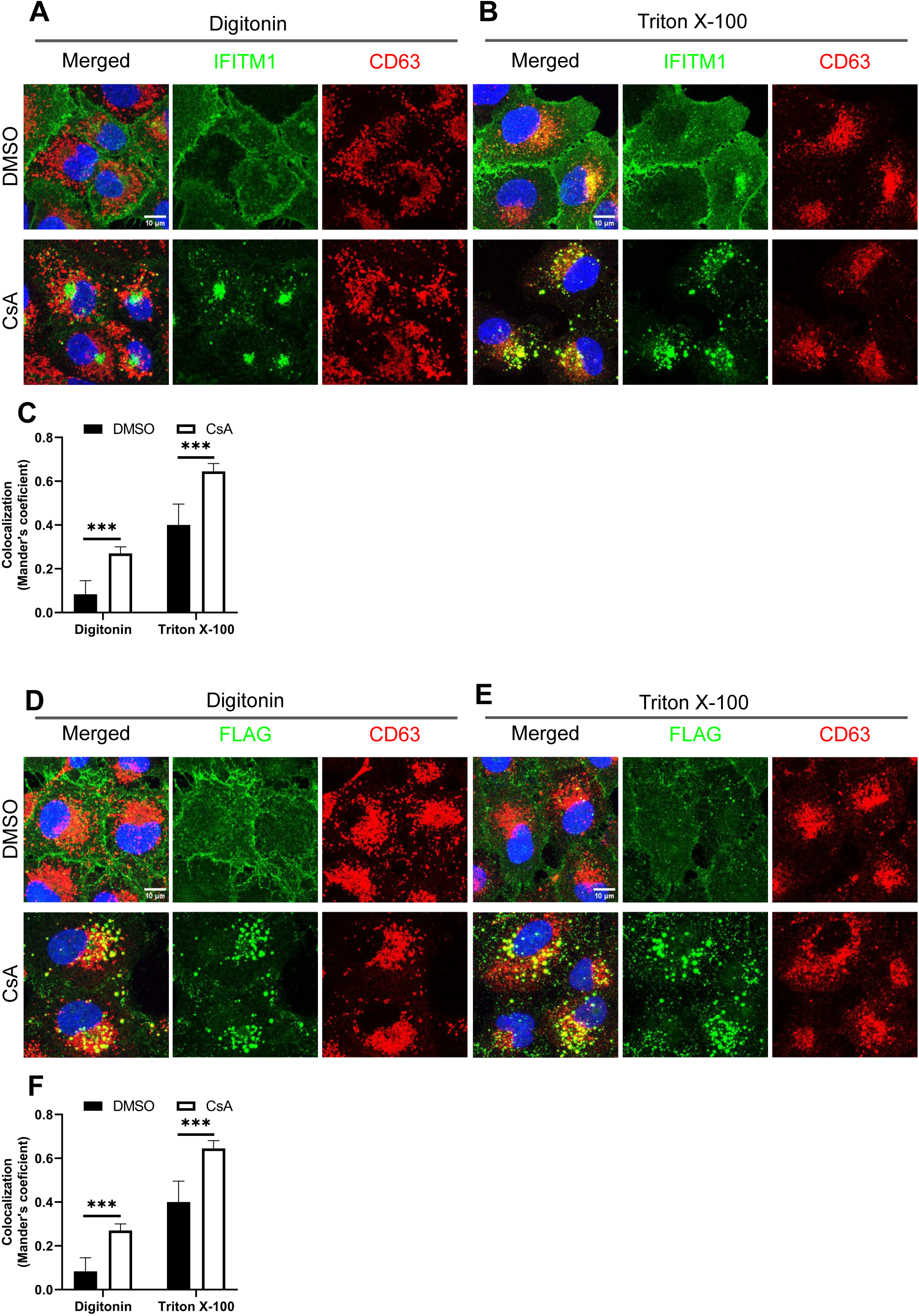
Disparate subcellular localizations of the IFITM1’s N- and C-termini following CsA-treatment. (A, B) A549.IFITM1-FLAG cells were fixed, permeabilized with digitonin (A) or TX-100 (B), and stained for IFITM1 and CD63. (D, E) As in panels A and B, but cells were stained for FLAG and CD63. (C, F) Colocalization of the IFITM1 N- or C-termini with CD63 under different conditions was determined for the maximum intensity projection images, using MOC.

**Figure S9.**
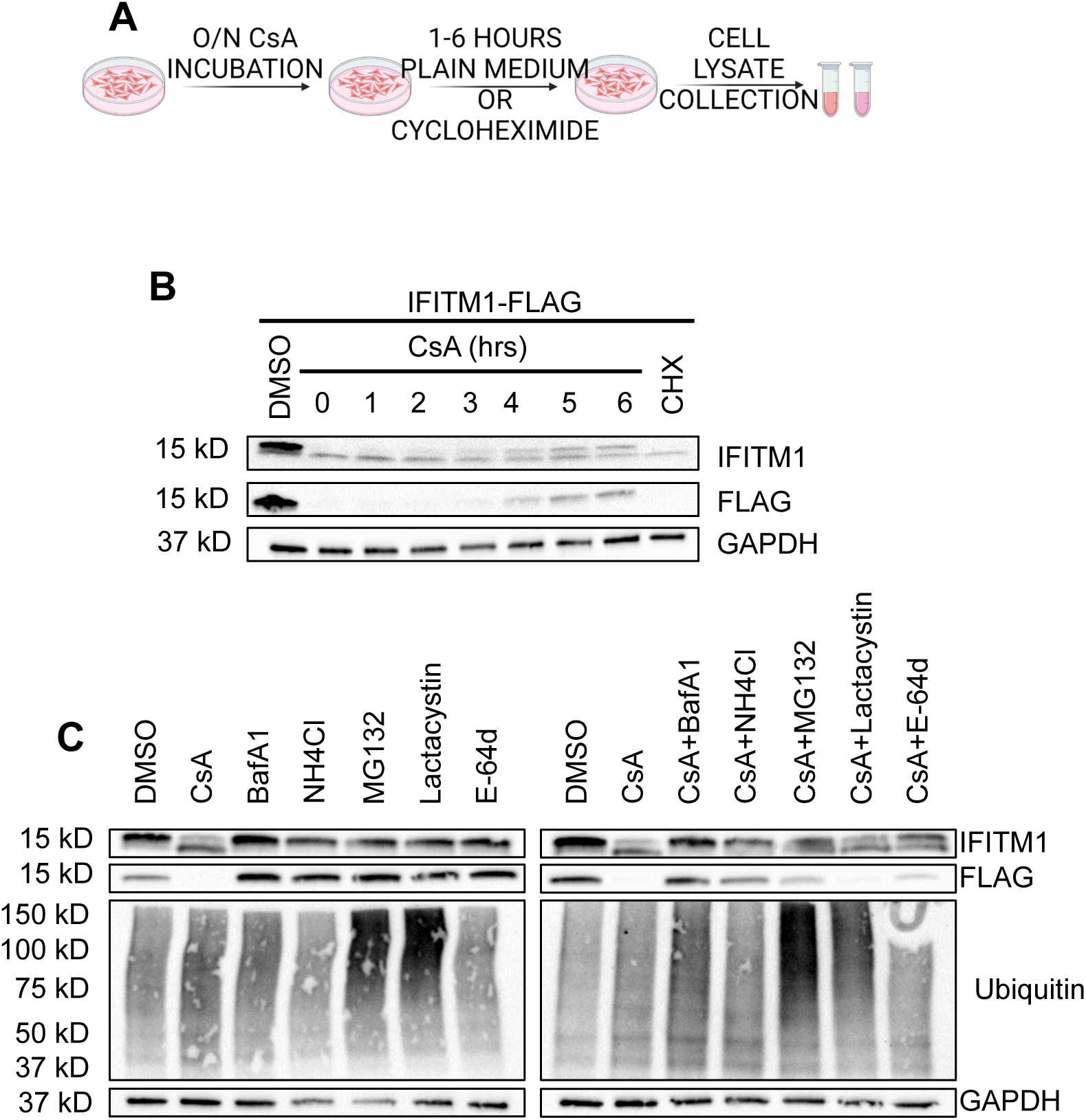
IFITM1 topology in CsA-treated cells is in line with the generally accepted type II topology. (A) Protocol schematics. Cells were kept with CsA in the medium overnight, washed, and incubated for the indicated times in a medium containing or lacking CHX (10 µg/mL). (B) A549.IFITM1-FLAG cells were treated as described in (A), harvested, and analyzed by Western blotting for IFITM1 and FLAG. (C) A549.IFITM1-FLAG cells were pre-incubated with inhibitors of endosome acidification, Bafilomycin A1 (BafA1, 1 μM) or ammonium chloride (NH_4_Cl, 40 mM), proteasomal inhibitors, MG132 (10 μM) or Lactacystin (10 μM), or the pan-cathepsin inhibitor, E-64d (20 μM). After one hour, CsA was added to the medium, cells were incubated for 6 more hours, harvested, lysed, and examined by Western blotting using anti-IFITM1, -FLAG, - Ubiquitin, or -GAPDH antibodies.

**Figure S10.**
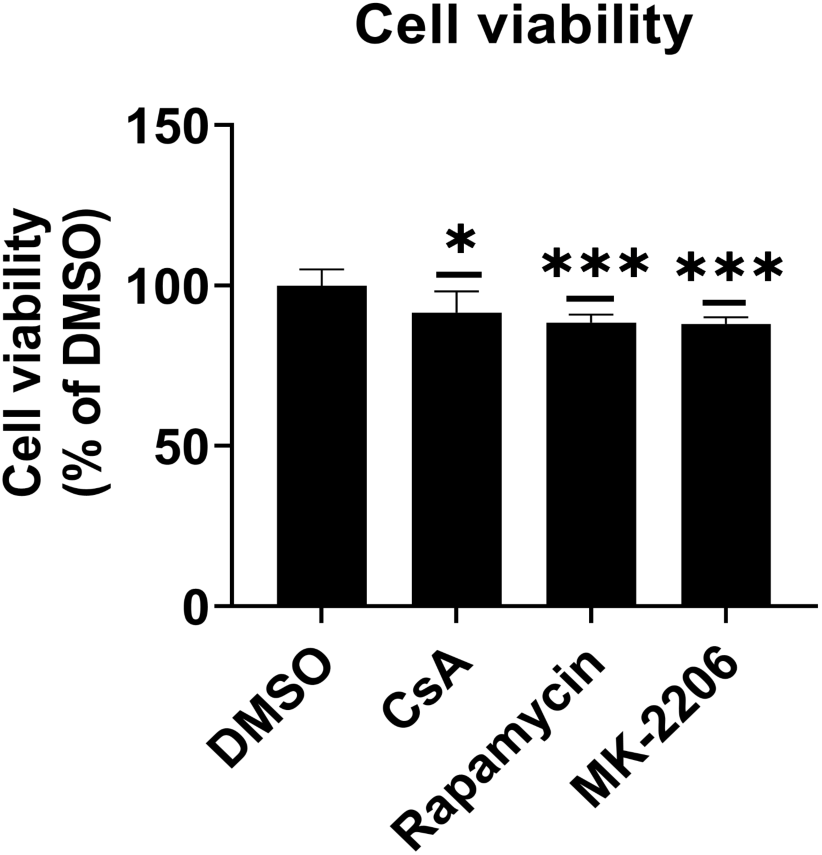
Effects of CsA, rapamycin, and MK-2206 treatment on A549 cell viability. Data represents cell viability of A549.vector cells treated with DMSO, CsA (20 µM), rapamycin (20 uM) or MK-2206 (10 µM) for 90 minutes. * p < 0.05; *** p < 0.001; ns, not significant. See also Figure 5.

**Figure S11.**
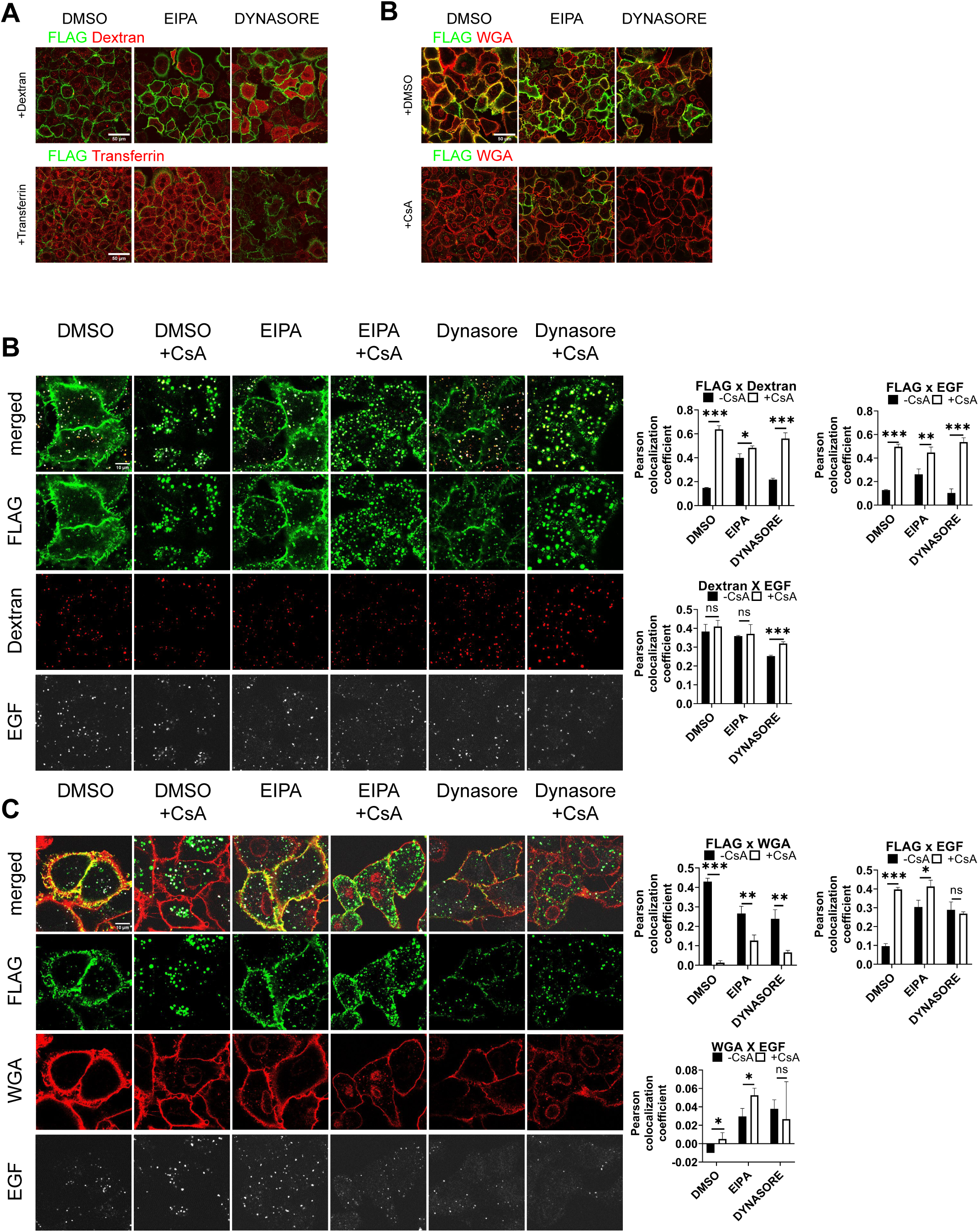
CsA induces redistribution of IFITM1 from the plasma membrane predominantly via dynamin-independent pathway. (A) A549.IFITM1-C-Flag cells were pre-treated with the respective compounds (EIPA 50 µM, Dynasore 120 µM) before exposure to cargo (EIPA for 30 min, Dynasore for 15 min, for details, see the Methods section) for the designated uptake pathway—Dextran (macropinocytosis) or transferrin (dynamin-dependent endocytosis)— to assess inhibition. The integrated intensity and colocalization with the plasma membrane marker, IFITM1-C-Flag, were quantified and plotted. Scale bar 10 µm. (B) A549.IFITM1-C-Flag cells were pre-treated with the appropriate inhibitor and incubated with fluorescently tagged EGF on ice. The medium was replaced with a fresh medium containing dextran and either CsA or DMSO, with inhibitors maintained throughout. After 30 minutes, cells were fixed and imaged. The colocalization of Flag (IFITM1) with the respective markers (dextran, EGF) and between the markers was analyzed, and integrated intensity was measured. Scale bar 10 µm. (C) A549.IFITM1-C-Flag cells were treated as described in (B), but dextran treatment was omitted. Instead, the plasma membrane was stained with WGA post-fixation. The colocalization of Flag (IFITM1) with the respective markers (WGA, EGF) and between the markers was analyzed, and integrated intensity was measured. For more details, see the Methods section.

**Figure S12.**
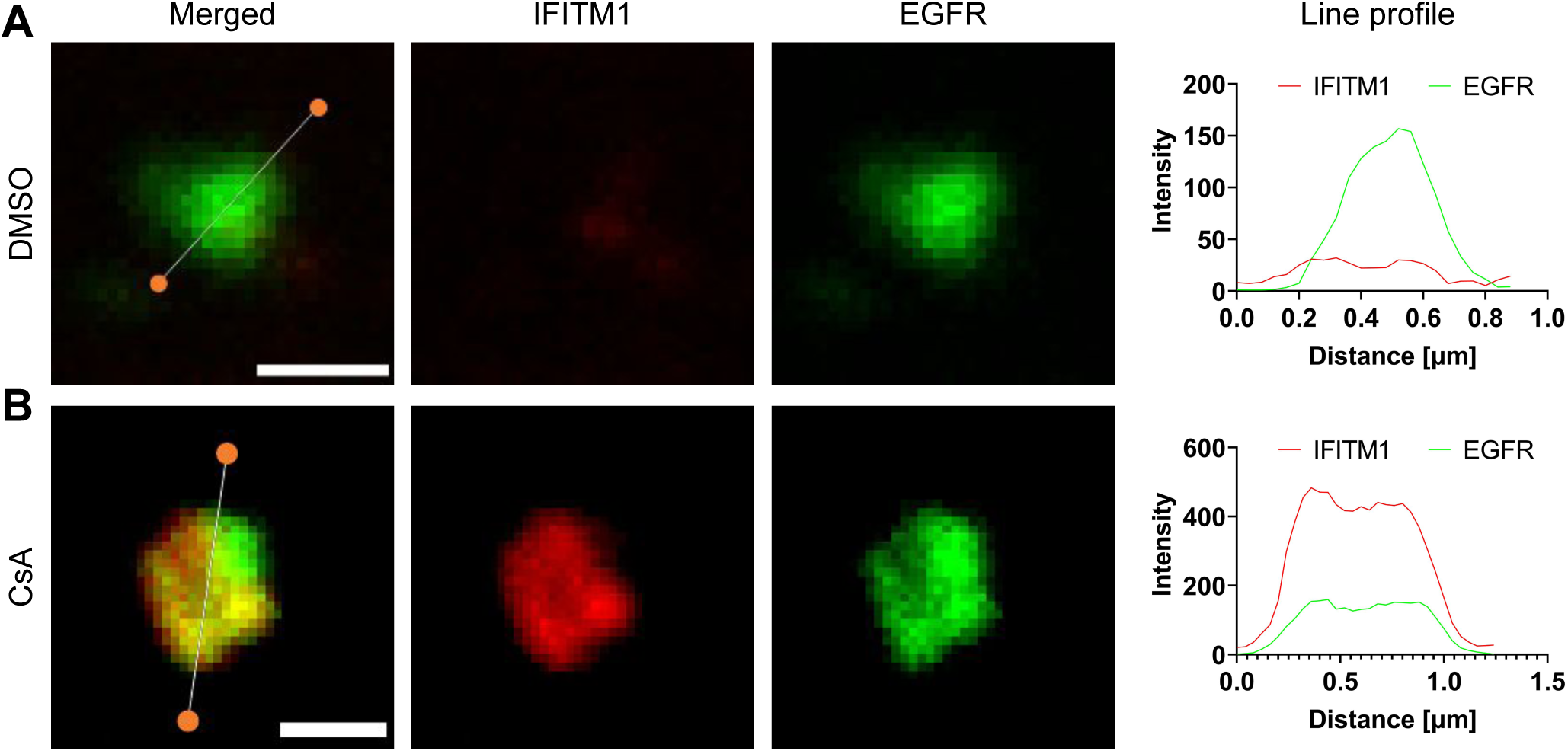
Redistribution of IFITM1 from the plasma membrane to the late endosome. A549.IFITM1-C-FLAG cells were treated with CHX, EGF and either DMSO (A) or CsA (B), as in Fig. 4B. Cells were fixed, permeabilized with TX-100, incubated with anti-IFITM1 and anti-EGFR antibodies, and stained with secondary antibodies conjugated to STED-compatible fluorophores, STAR RED and STAR 580. Representative images of n>20 analyzed endosomes are shown. Line histograms for selected endosomes are shown. Scale bar is 0.5 µm.

**Figure S13.**
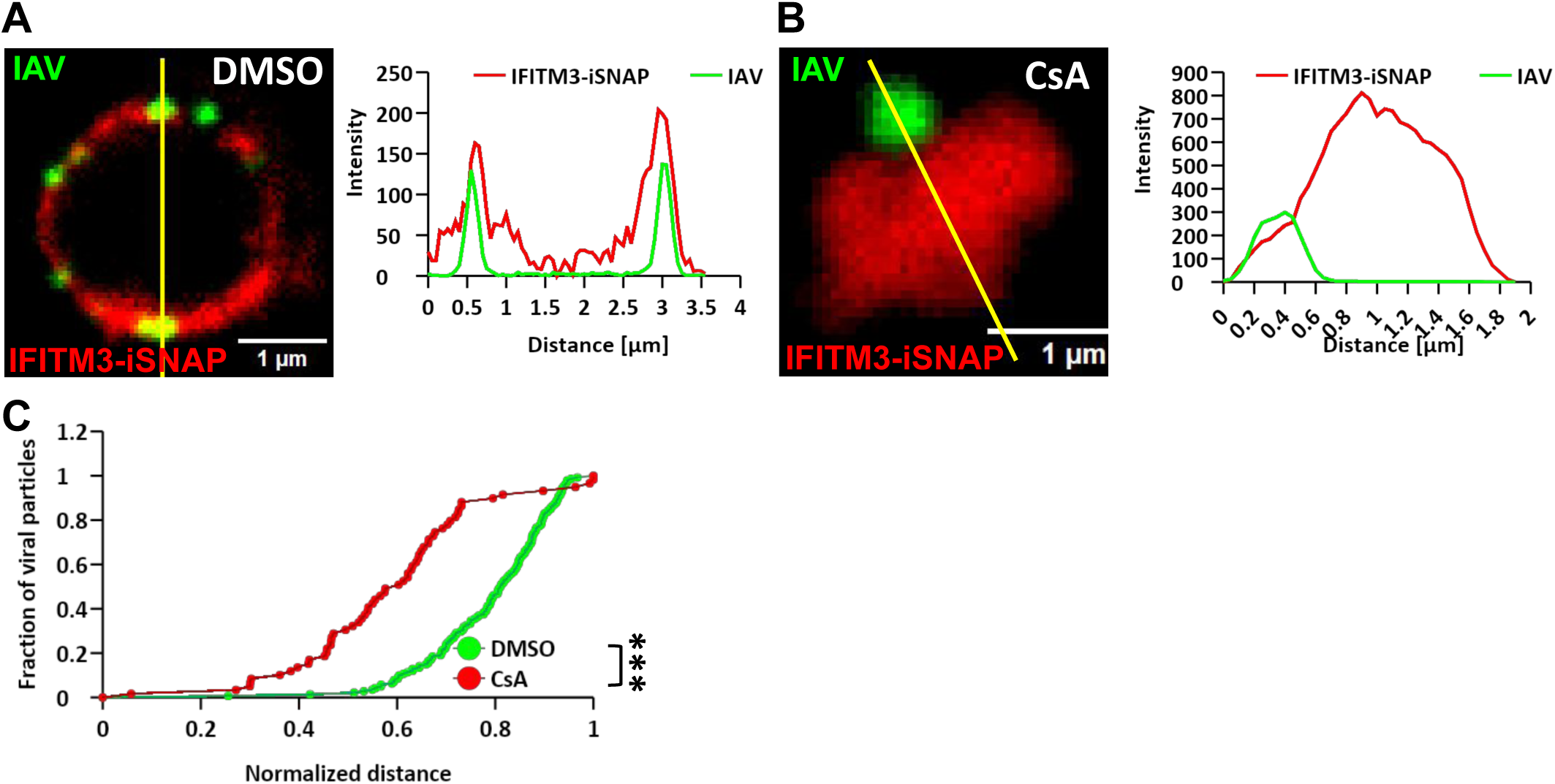
CsA induces redistribution of IFITM3 to the interior of late endosomes. (A-C) A549.IFITM3-iSNAP cells were pre-incubated with DMSO (A) or 20 µM of CsA (B) for 1.5 hours and spin-infected with AF-568 labeled IAV at MOI of 2. Infection was allowed to proceed for 1 hour in the presence of DMSO or CsA, at which time, cells were stained with SNAP-Cell 647-SiR for 30 min, washed and incubated with fresh medium for additional 30 min to remove unbound dye. Cells were fixed and imaged using STED super-resolution microscopy. Right graphs in panels A and B show the line intensity profiles across the endosomes and IAV particles corresponding to images on the left. (C) The distance of individual IAV particles to the center of the endosome was measured and normalized to the endosome’s radius. Distances for IAV from at least 5 endosomes were measured and plotted for each condition. Lines and bars are means and S.D. ***, p < 0.001.

**Movie S1. Staining of IFITM1 in the presence of DMSO.** A549.IFITM1-C-FLAG cells were stained with anti-Flag antibody conjugated with AF-647 to visualize IFITM1 (green) and Hoechst to visualize nuclei (blue) and imaged in the presence of DMSO (vol %?) for indicated time. Time is in a mm:ss format. Movie is related to Figure S11.

**Movie S2. Staining of IFITM1 in the presence of CsA.** A549.IFITM1-C-FLAG cells were stained with anti-Flag antibody conjugated with AF-647 to visualize IFITM1 (green) and Hoechst to visualize nuclei (blue) and imaged in the presence of 25 µM CsA for indicated time. Time is in a mm:ss format. Movie is related to Figure S11.

## Notes

### Competing Interest Statement

The authors have declared no competing interest.

